# Mechanistic Insight into the Conformational Changes of Cas8 upon Binding to Different PAM Sequences in the Transposon-Encoded Type I-F CRISPR-Cas System

**DOI:** 10.1101/2024.02.01.578494

**Authors:** Amnah Alalmaie, Raed Khashan

**Author notes:** Corresponding author: Raed Khashan, PhD Associate Professor Pharmaceutical Sciences Division of Pharmaceutical Sciences Long Island University Brooklyn, NY 11201.

## Abstract

The INTEGRATE system is a gene-editing approach that offers advantages over the widely used CRISPR-Cas9 system. It does not introduce double strand breaks in the target DNA but rather integrates the desired DNA sequence directly into it. The first step in the integration process is PAM recognition, which is critical to understanding and optimizing the system. Experimental testing revealed varying integration efficiencies of different PAM mutants, and computational simulations were carried out to gain mechanistic insight into the conformational changes of Cas8 during PAM recognition. Our results showed that the interaction between Arg246 and Guanine at position (-1) of the target strand is critical for PAM recognition. We found that unfavorable interactions in the 5’-AC-3’ PAM mutant disrupted this interaction and may be responsible for its 0% integration efficiency. Additionally, we discovered that PAM sequences not only initiate the integration process but also regulate it through an allosteric mechanism that connects the N-terminal domain and the helical bundle of Cas8. This allosteric regulation was present in all PAMs tested, even those with lower integration efficiencies, such as 5’-TC-3’ and 5’-AC-3’. We identified the Cas8 residues that are involved in this regulation. Our findings provide valuable insights into PAM recognition mechanisms in the INTEGRATE system and can help improve the gene-editing technology.

## Introduction

CRISPR-Cas systems are adaptive immune systems found in archaea and bacteria [1] The main components of the CRISPR-Cas system are the CRISPR repeat-spacer array and Cas genes encoding Cas proteins. Generally, Cas proteins drive CRISPR immunity in three distinct molecular stages, which are shared between major CRISPR-Cas systems: the first is the spacer acquisition stage, also known as “adaptation,” the second is the pre-crRNA processing, finally is the interference stage [1]. At the spacer acquisition stage, Cas1 and Cas2, common in most CRISPR-Cas systems, seize a segment of the target DNA “protospacer” and insert it at the 5 ′ ends of a CRISPR array. In the second stage, the pre-crRNA processing step, the CRISPR array is transcribed into a long immature crRNA known as “pre-crRNA,” which will eventually be processed into mature, small crRNAs Next is the interference stage; the mature crRNA bound by Cas proteins scans the DNA for a <u>P</u>rotospacer <u>A</u>djacent <u>M</u>otif (PAM) sequence; once the PAM sequence is located, the base pairing between the crRNA spacer and the complementary DNA protospacer takes place, followed by subsequent cleavage by a dedicated nuclease domain [1].

CRISPR-Cas systems are generally classified into two main classes and six types [1,2]. Class 1 utilizes a multi-subunit crRNA–effector complex consisting of several Cas proteins bound together with the mature crRNA Class 1 comprises three types (type I, III, and IV) and 12 subtypes [3]. Type I CRISPR-Cas system is the most predominant and diverse group of CRISPR-Cas systems, accounting for ∼90 % of identified systems [1]. The nucleases responsible for target cleavage are Cas3 in type I and Cas10 in type III [1–3]. Type IV has no adaptation and nuclease genes, and the mechanism of action is unknown [3]. Class 2 CRISPR-Cas system is more straightforward, in which one protein realizes all functions of the multiprotein effector complex [1]. There are three types: type II, type V, and type VI, with Cas9, Cas12, and Cas13, respectively, that fulfill all the functions of the multiprotein effector complex [1,3].

The presence of short CRISPR arrays similar to those found in nuclease-deficient type I-B, type I-F, and type V-K CRISPR-Cas systems has been reported in several bacterial Tn7-like transposons [3]. In these transposons encoded CRISPR-Cas systems, Cas1, and Cas2 genes, which are responsible for spacer acquisition, are absent. In addition to the absence of the nucleases Cas3 and Cas12 genes, which are responsible for target DNA cleavage [4–8] . It has been shown that the transposition mechanism of the *Vibrio cholera* Tn6677 transposon (VcTn6677) is based on a synergy between the transposition components and a nuclease-deficient type I-F CRISPR-Cas system *Vibrio Cholera* Tn6677 transposon consists of two main components; the Cascade subunits, which consist of Cas6, six subunits of Cas7, and fused Cas8/5 (simply Cas8), and the transposition subunits (TnsA, TnsB, TnsC, and TniQ) [4–8]. The whole complex is called the INTEGRATE system (<u>IN</u>sert <u>T</u>ransposable <u>E</u>lements by <u>G</u>uide <u>R</u>NA-<u>A</u>ssisted <u>T</u>ar<u>gE</u>ting) [4–8]. Interestingly, the TniQ gene (TnsD-like) is present within the Cas operon and not within the Tns operon, **as shown in Fig. 1** [5].

**Fig. 1:**
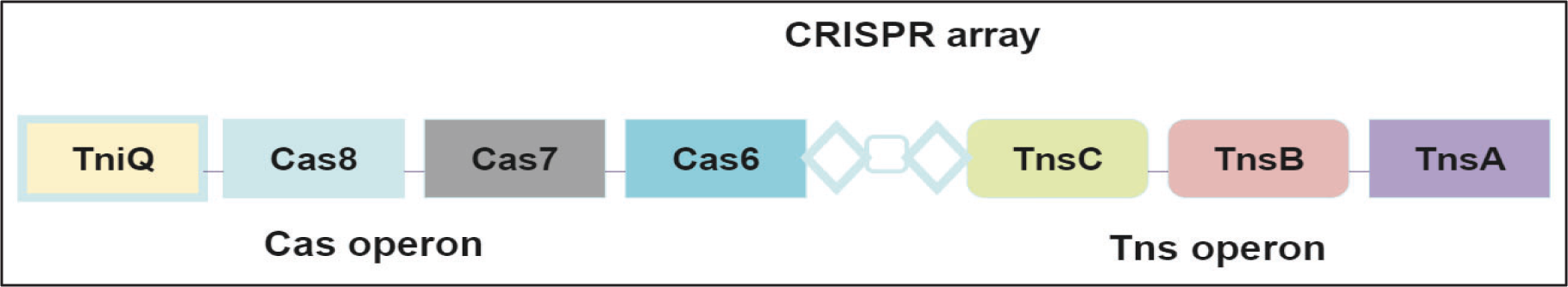
Overall architecture of the V. cholerae TniQ–Cascade complex [5] Created with BioRender.com

DNA binding occurs in three significant steps, confirmed by independent studies [5,7–10]. Firstly, the Cascade components assemble around CRISPR-RNA (crRNA), followed by the Cascade/crRNA complex binding to the TniQ protein [5,7,8]. The third step involves scanning the target DNA to find a <u>P</u>rotospacer <u>A</u>djacent <u>M</u>otif (PAM) sequence; the PAM recognition initiates a local unwinding of a double-stranded DNA, followed by a base pairing between the nucleotides region of the spacer and the complementary protospacer at the first eight PAM-proximal nucleotides of the crRNA (termed seed sequence) [5,7,8]. This initial base-pairing is extended between the spacer and protospacer, resulting in a complete hybridization between the target strand and the crRNA while displacing the non-target stand to generate what is known as the R-loop structure, which in turn initiates the transposition process, **as shown in Fig. 2** [5,7,8].

**Fig. 2:**
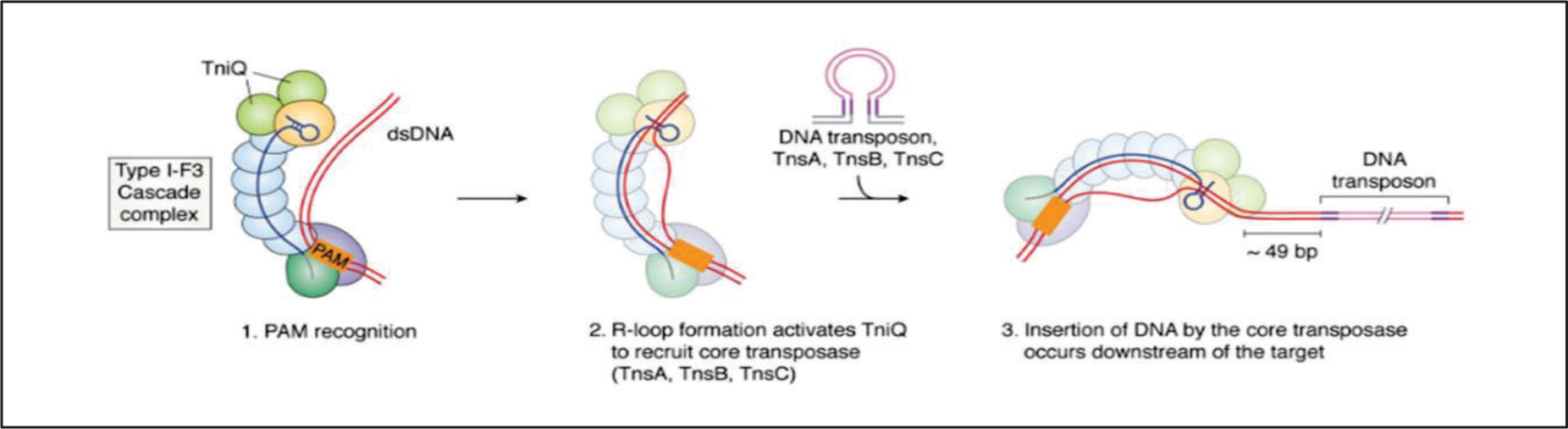
The RNA-guided DNA-transposition process. [11].

In the type II CRISPR-Cas system, the *Streptococcus Pyogenes* Cas9 nucleases recognize a PAM sequence of 5′-NGG-3′ located in the major groove of the target DNA Cas9 consists of two major lobes, a recognition (REC) and a nuclease (NUC) lobe [12,13] . PAM recognition involves a specific interaction between the arginine residues located in the PAM interacting domain with the first and second guanine bases of the PAM sequence. The NUC lobe contains the two nuclease domains that are required for DNA cleavage (RuvC and HNH), in addition to the PAM interacting (PI) domain, **as shown in Fig. 3** [12,13].

**Fig. 3:**
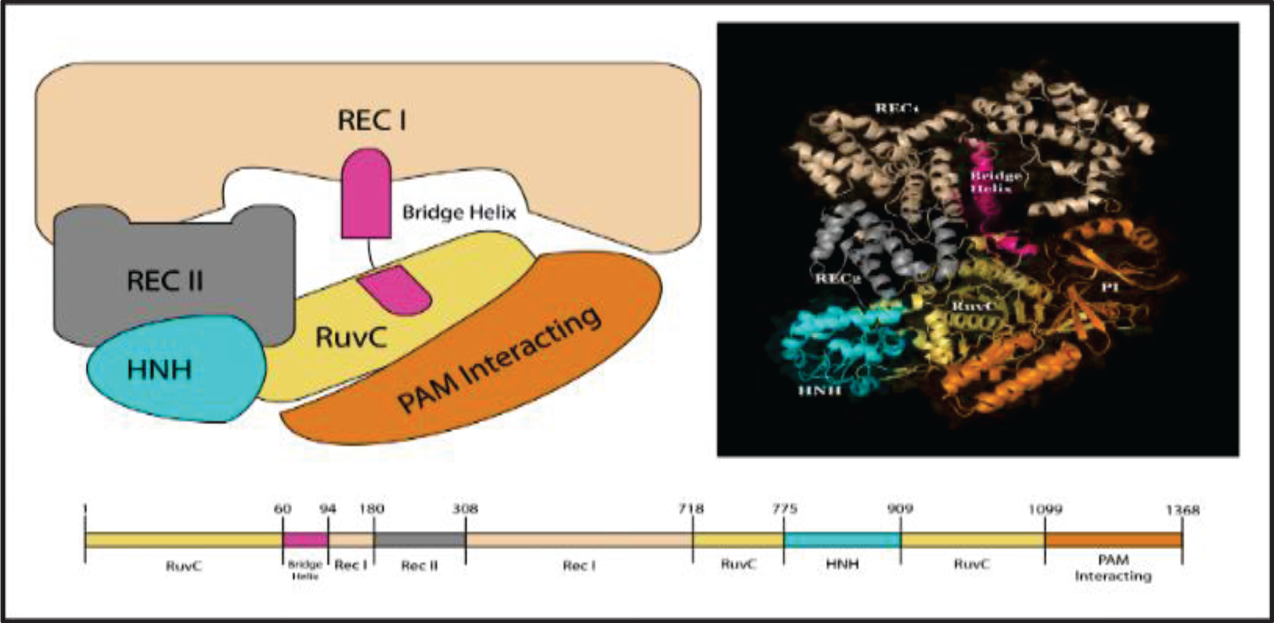
Cas9 Protein domains. The Cas9 protein comprises six domains: Rec I, Rec II, Bridge Helix, RuvC, HNH, and PAM Interacting. (Crystal image rendered from PDB code: 4CMP. [12].

In the type I CRISPR-Cas system, the PAM recognition is promiscuous due to the minor groove non-sequence specific contact between the PAM sequence and Cas8 protein [14]. The DNA’s minor groove can deform and change its geometry to accommodate the protein bound to it [15]. This inherent flexibility of the minor groove results in a diverse combination of different PAMs recognized with varying affinities . In 2019, Sternberg’s group tested different dinucleotide PAM sequences to examine each sequence’s integration efficiency in the INTEGRATE system [4]. The integration efficiency was measured for mutant 5′-CN-3′ PAM systems. All mutants with 5′-CN-3′ PAM show indistinguishable integration efficiency compared to the wild-type 5′-CC-3′ PAM, indicating the unique behavior of Cytosine at the -2 position (again, 5’-CN-3’) [4].

On the other hand, all PAM sequences with mutations in the -1 position (5′-NC-3′) show less integration compared to the wild-type PAM [4] . Interestingly, the 5′-AC-3′ PAM sequence shows no integration efficiency[4]. The ability of Cas8 to recognize different PAMs is a clear indication of PAM recognition flexibility in the INTEGRATE system, in contrast to the strict requirements to identify PAM sequences by Cas9 enzyme in Type II CRISPR systems.

The target DNA-bound V. Cholerae TniQ-Cascade complex’s cryo-EM structure reveals critical structural features of the Cas8 protein upon binding to the 5′-CC-3′ PAM sequence Cas8 protein consists of three domains, the N-terminal domain (Cas8NTD, aa 1-275), the middle helical bundle (Cas8HB, aa 276-384), and the C-terminal domain (Cas8CTD, aa 385-640) [8]. The N-terminal domain of Cas8 is directly involved in PAM recognition. In contrast, the Cas8 helical bundle (Cas8HB) does not participate in the PAM recognition because its location is distal. Instead, Cas8HB is expected to trigger the activation of TniQ to initiate the DNA transposition process [8].

In the NTD of Cas8, the arginine residue (Arg246) forms a stacking interaction with the guanine nucleotide on the target strand, allowing the arginine residue to act as a wedge to separate the double-stranded DNA [5]. The separation of the double-stranded DNA allows crRNA to invade and base pairs with the target strand while displacing the non-target strand to form the R-loop Structure [6,7]. Wedge insertion is structurally conserved across different Type I systems, but sequence variability enables the recognition of varying PAM sequences [1,6,10]. Much progress is needed in the INTEGRATE system to understand the structural features of Cas8 involved in PAM recognition.

Many protein structures have been resolved by one of three main techniques: X-ray crystallography (X-Ray), nuclear magnetic resonance (NMR), and cryo-electron microscopy (Cryo-EM); however, these techniques provide static structures. To understand the biological function of these proteins, we must understand the motion of the atoms. Molecular dynamics simulation (MD) complements conventional experiments; MD enables researchers to investigate phenomena that cannot be studied experimentally [16,17]. We have carried out MD simulations to study the molecular details of the conformational changes in Cas8 upon binding to the wild-type PAM (5’-CC-3’) and mutated PAM sequences (5’-CG-3’, 5’-TC-3’ and 5’-AC-3’), **as shown in** Fig. 4. To the best of our knowledge, this work is the first effort to use MD simulation to understand the conformational changes in the INTEGRATE system.

**Fig. 4.**
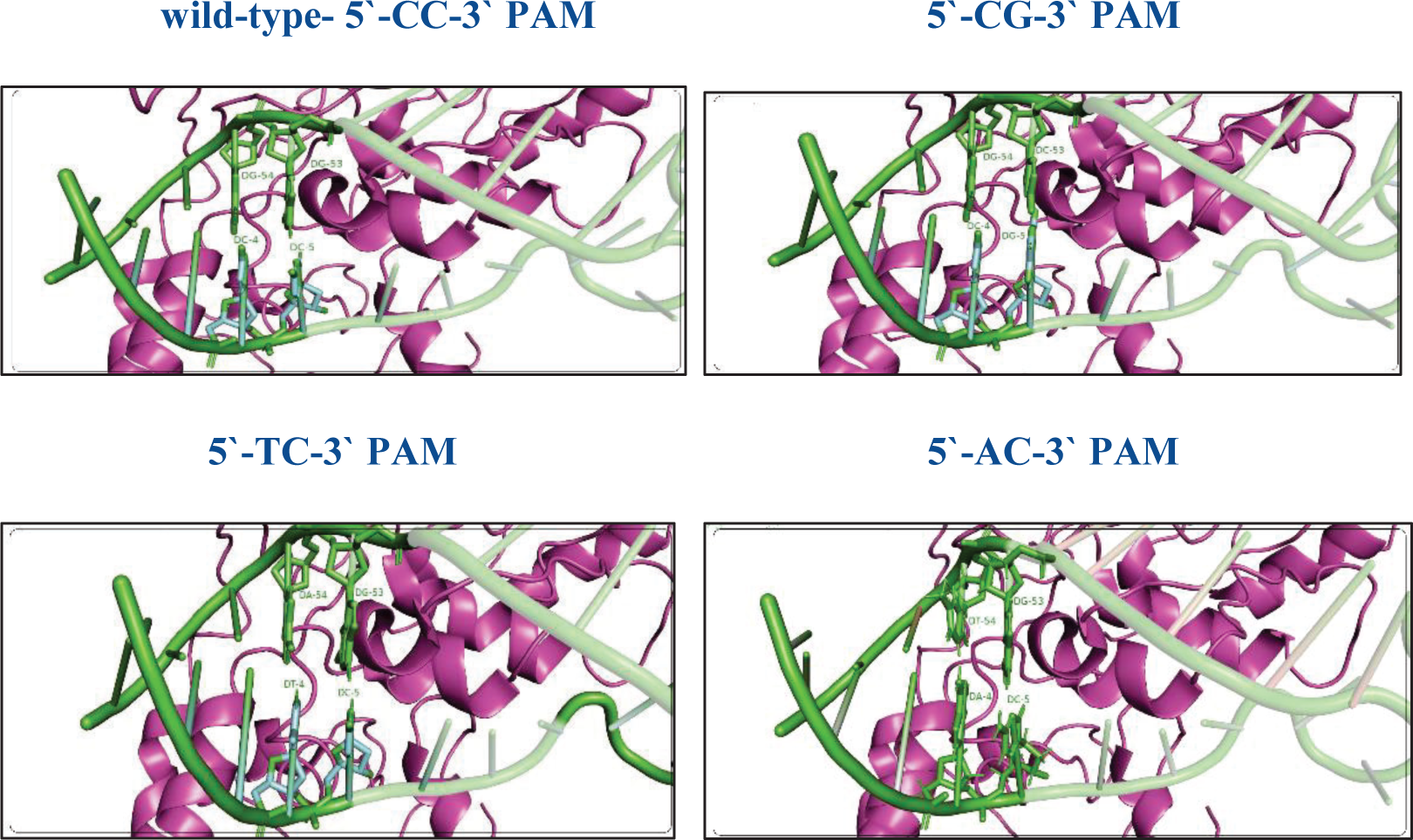
**Schematic representation of the present work**

The results provide mechanistic insights into how mutating the PAM sequence influences its recognition and the underlying effect of such mutations on the dynamic and interactions with the Cas8 subunit. The key nonbonded contacts have been explored comprehensively to understand the impact of the mutation on these critical interactions. The results provide deep insight into the mechanism of PAM recognition in the INTEGRATE system at an atomistic level which could be useful in the engineering of customized CRISPR-Cas systems and expand the targeting space.

## Method

### Structure Models

Cryo-EM structure is available in the RCSB-PDB database of the ternary TniQ-Cascade complex bound to crRNA, and DNA (PDB code:6PIJ) containing 5′-CC-3′ PAM sequence was used as a starting point for the MD simulation [5]. The structure can be considered an intermediate state due to an incomplete R-loop structure, in which an apparent density of 22 nucleotides of the target strand within the 32-bp DNA protospacer was observed. Regarding the non-target strand, there is a clear density of only eight nucleotides; the remaining nucleotides were modeled based on the non-target strand of the type I-E CRISPR-Cas system (PDB code: 5U0A) [18]. The double-stranded region was modeled using web3DNA [19] . The missing protein residues in the structures were modeled with two software (modeler and Charmm-Gui) [20].

Three models have been built based on the structure with a PDB code:6PIJ, PAM mutant models 5′-CG-3′, 5′-TC-3′, and 5′-AC-3′. Two of them (5′-AC-3′, 5′-TC-3′ PAM mutant models) have a mutation at the -2 position (5′-NC-3′), where this position severely affects the integration efficiency [4]. The 5′-AC-3′ PAM mutant model was built by mutating Cytosine to Thymine at position -2 of the non-target strand and Guanine to Adenine at position -2 of the target strand. In contrast, the 5′-TC-3′ model was prepared by mutating Cytosine at position -2 to Adenine and Guanine at position -2 to Thymine. These mutations were chosen to be consistent with the experimental results for which the integration efficiency was mustered [4]. For the 5′-TC-3′ PAM, the experimental results indicated 60% integration efficiency, whereas the 5′-AC-3′ PAM completely inhibits the integration [4]. Regarding the third model, the PAM mutation was introduced at the -1 position (5′-CN-3′), Cytosine at position -1 of the non-target strand was mutated to Guanine, and the Guanine at position -1 of the target strand was mutated to Cytosine to build 5′-CG-3′ PAM model. All mutations were introduced using web3DNA [19] . The numbering of the PAM proximal region, including the di-nt PAM sequence, **is shown in Fig. 5**

**Fig. 5.**
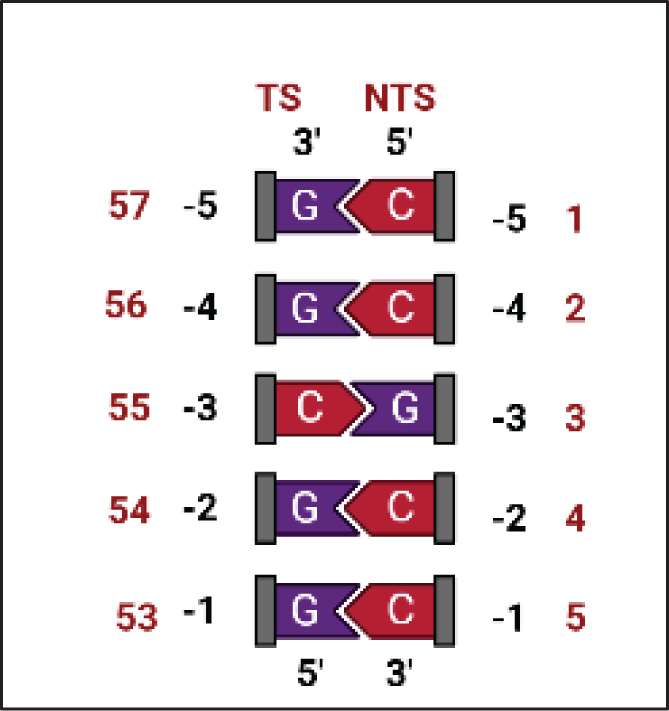
Schematic drawing of the PAM proximal region. The numbers in red are based on the numbering of PDB code: 6PIJ The di-nt PAM sequence is labeled as 4 and 5 of the TS, 54 and 53 of the NTS. Created with BioRender.com

### Simulation Method

A popular molecular dynamic package (GROMACS) was used for setting up, running, and analyzing the MD simulations [21]. We carried out four independent simulations for a total of 4 μs; each model contains a different PAM sequence, 5′-CC-3′, 5′-CG-3′, 5′-TC-3′, and 5′-AC-3′, respectively. All models were prepared and simulated with identical parameters. Amber ff12SB force field was applied for the protein and nucleic acids, including f99bsc0_chiOL3 modification for RNA and ff99bsc0 modification for DNA [22] . This force field has been shown to adequately describe the conformational dynamics of CRISPR MD simulations. The complex was solvated in a triclinic box in each model, and a TIP3P water model was used to simulate water. The simulation box was charge-neutralized by adding Na+ counterions. Each model has ∼ 474453 atoms, including water molecules and ions. Energy minimization was used to remove clashes and bad contacts that would result in an unstable simulation. Fifty thousand (50000) steps of the steepest- descents energy minimization algorithm were used.

Following the energy minimization stage, two equilibrating simulations were run to bring the system temperature and pressure to the desired values. NVT (constant <u>N</u>umber of atoms, <u>V</u>olume, and <u>T</u>emperature) equilibration and NPT (constant <u>N</u>umber of atoms, <u>P</u>ressure, and <u>T</u>emperature) equilibration. In NVT equilibration, the system was heated up from 0 to 300 K in the canonical ensemble (NVT) by running four NVT simulations, **as shown in Table 1**. Temperature control (300 K) was performed by coupling the system to a Nose-Hover thermostat. NPT equilibration was performed for 20 ns. Pressure control was performed by coupling the system to a Parrinello-Rahman barostat at a reference pressure of 1 atm.

**Table 1:** The condition of the four NVT simulations used for the project.

| **NVT Simulation Temperature** | **NVT Simulation Time** | **Position Restraints** |
| --- | --- | --- |
| 0K | 20ps | 300 kcal-mol |
| 0-100K | 50ps | 100 kcal-mol |
| 100-200K | 100ps | 25 kcal-mol |
| 200-300K | 200ps | None (unrestrained) |

An integration time step of 2 fs was used for the production run to carry out MD simulation. For each model, the simulation was carried out in the NPT ensemble for 1 μs of production runs. The Particle-Mesh Ewald (PME) summation method was used to compute the long-range electrostatic interactions, with Fourier spacing of 1.2 nm and fourth-order interpolation. The force-switch method was used to treat the nonbonded interactions (van der Waals) with a cutoff of 1.2 nm. The LINCS algorithm constrained all the hydrogen atoms bonded to the protein-heavy atoms.

## Result

### Structural Evaluation

GROMACS utilities were used to evaluate the structural parameters such as RMSD, RMSF, and the number of hydrogen bonds formed between Cas8-NTD and PAM-proximal sequence with the donor-acceptor set at a cutoff of 0.35 nm. All 2D plots were generated using Grace. The root means square deviation (RMSD) of the backbone atoms of the whole complex and Cas8 subunit was evaluated to estimate the initial stability of the simulations over time, considering the first frame and the average frame as reference structures, **as shown in Fig. 6-8**

**Fig. 6:**
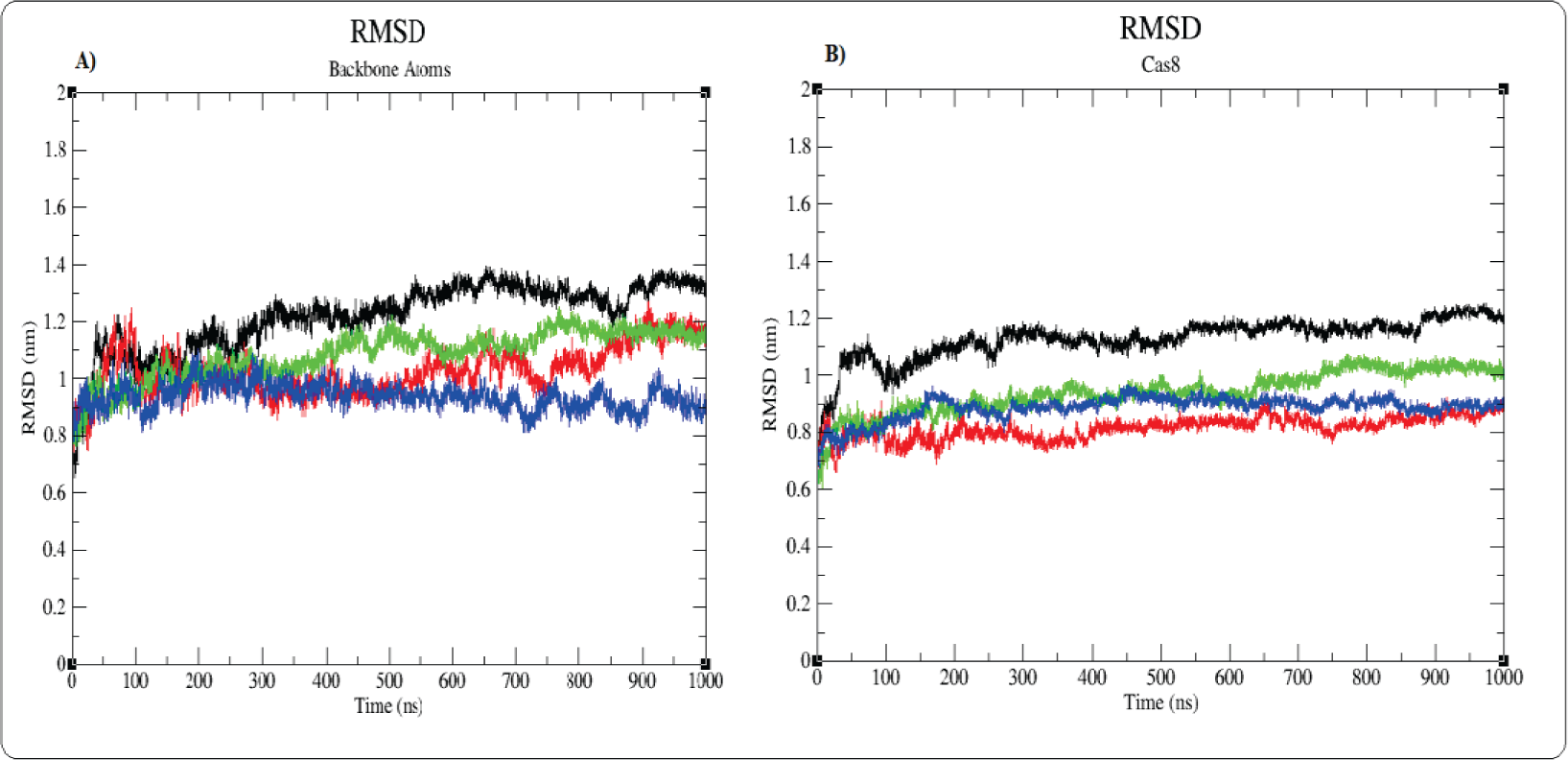
A) Backbone root-mean-square deviation (RMSD) compared to the first frame over the timescale of 1 μs. B) Cas8 root-mean-square deviation (RMSD) compared to the first frame over the timescale of 1 μs. *Black is 5′-CC-3′, blue is 5′-CG-3′, green is 5′-TC-3′, and red is 5′-AC-3′*

At the early stage of the simulation, the RMSD of the starting configuration of the wild- type 5′-CC-3′ PAM model experienced more increase compared to all other models, in which the average RMSD is 1.2 nm. The backbone of the 5′-CC-3′ PAM model reached a plateau at a higher value compared to the PAM-mutated models. The 5′-CG-3′ and 5′-TC-3′ PAM models displayed RMSDs of 0.94 nm and 1.09 nm, respectively. However, the backbone of the 5′-AC-3′ PAM model experienced some fluctuation throughout the trajectory, with an average RMSD of 1.3 nm. The RMSD values of the Cas8 indicated stability for the three systems; however, the wild type 5′-CC- 3′ PAM stabilized at higher values compared to other models. Overall, the backbone RMSD of all systems indicates that all PAM models maintained their structural integrity throughout MD simulations.

The RMSD values plateau at a relatively high value. The RMSD values with respect to the average structure are likely to offer a better perspective of the evolution of structural changes over the simulation. The backbone-RMSDs were found to be 0.29, 0.27, 0.28, and 0.29 nm, respectively, for wild-type 5′-CC-3′, 5′-CG-3′, 5′-TC-3′, and 5′-AC-3′, PAM systems. The backbone RMSD of the 5′-CG-3′ and 5′-AC-3′ showed more fluctuation compared to the wild- type 5′-CC-3′ PAM model, **as shown in Fig. 7**

**Fig. 7:**
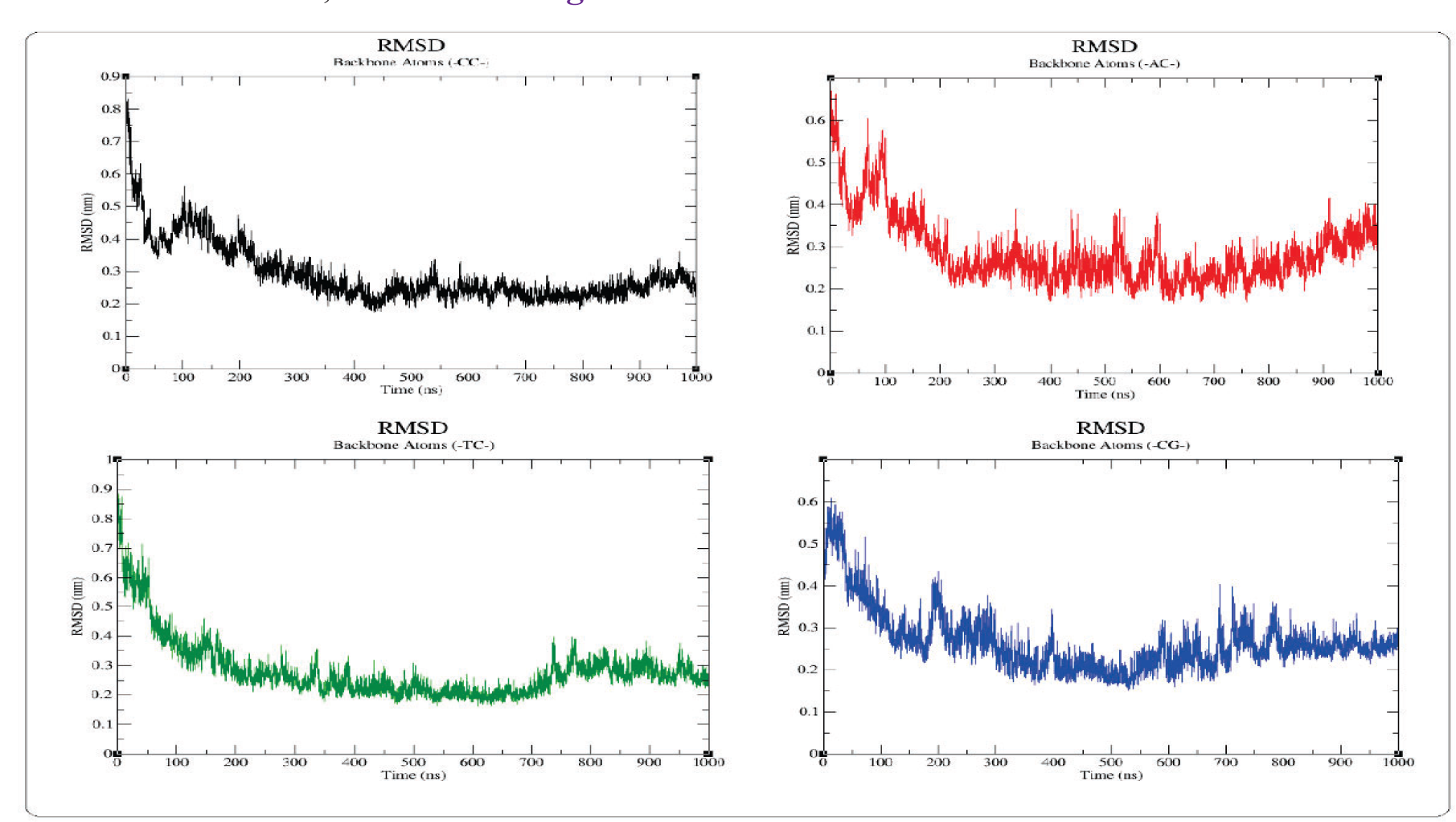
**Backbone RMSD values compared to the average structure over 1 μs simulation.**

**Fig. 8:**
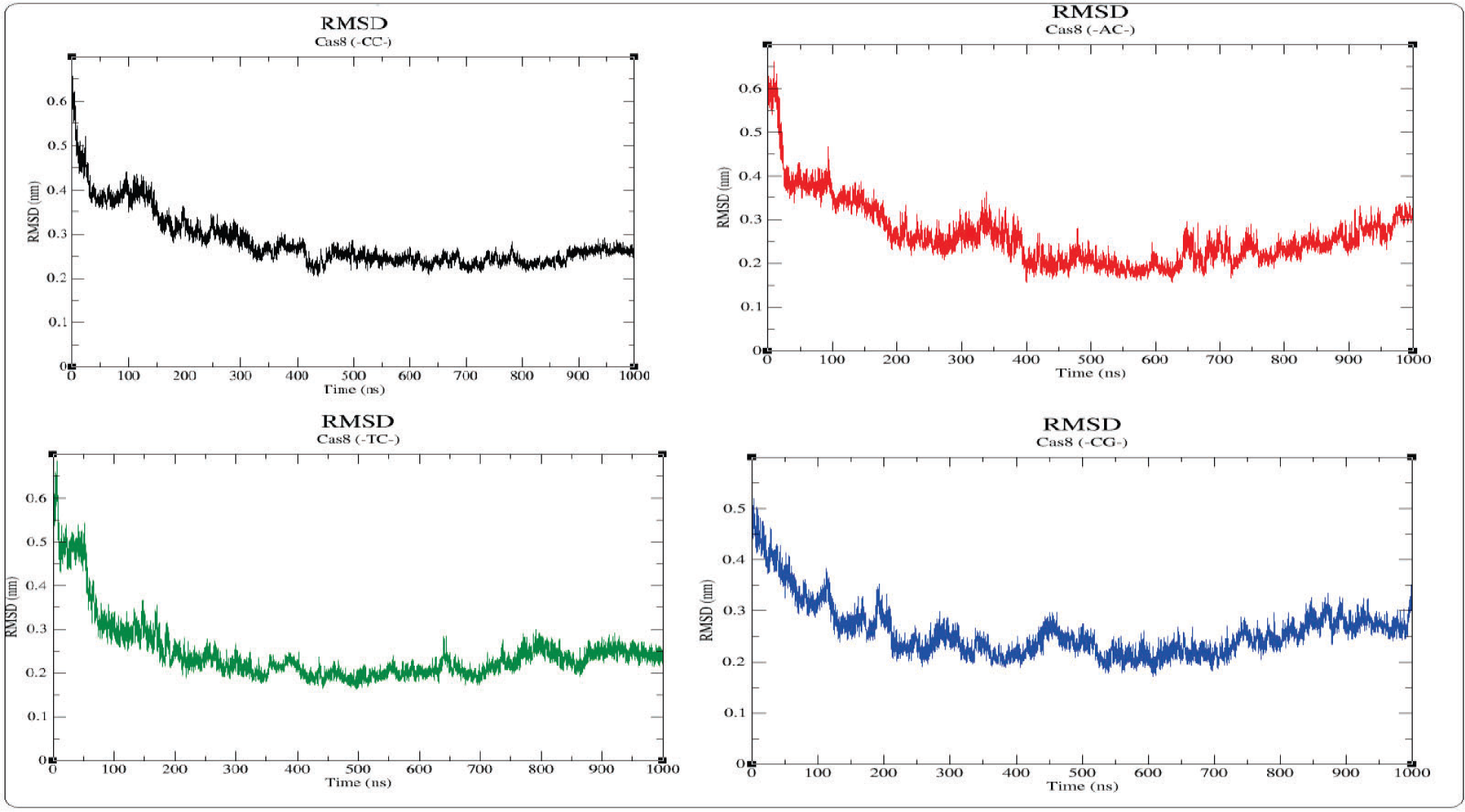
Cas8 root-mean-square deviation (RMSD) compared to the average frame over the timescale of 1 μs. *Black is 5′-CC-3′, red is 5′-AC-3′, green is 5′-TC-3′, blue is 5′-CG-3′*.

The RMSD values of Cas8 with respect to the average structure **are shown in Fig. 8**. The RMSD values revealed high stability for all models, wild-type-PAM was found to be stabilized after ∼ 300 ns, a similar trend was observed for the 5′-TC-3′ PAM. In contrast, the 5′-CG-3′ PAM model revealed lower stability. The mutant 5′-AC-3′ PAM model lingered on till 650 ns to reach a stable conformation to some extent.

The root-mean-square fluctuation (RMSF), which indicates how mobile each residue of Cas8 is around its mean position, was employed to understand the mobility and flexibility of Cas8 in each system, **as shown in Fig. 9**

**Fig. 9:**
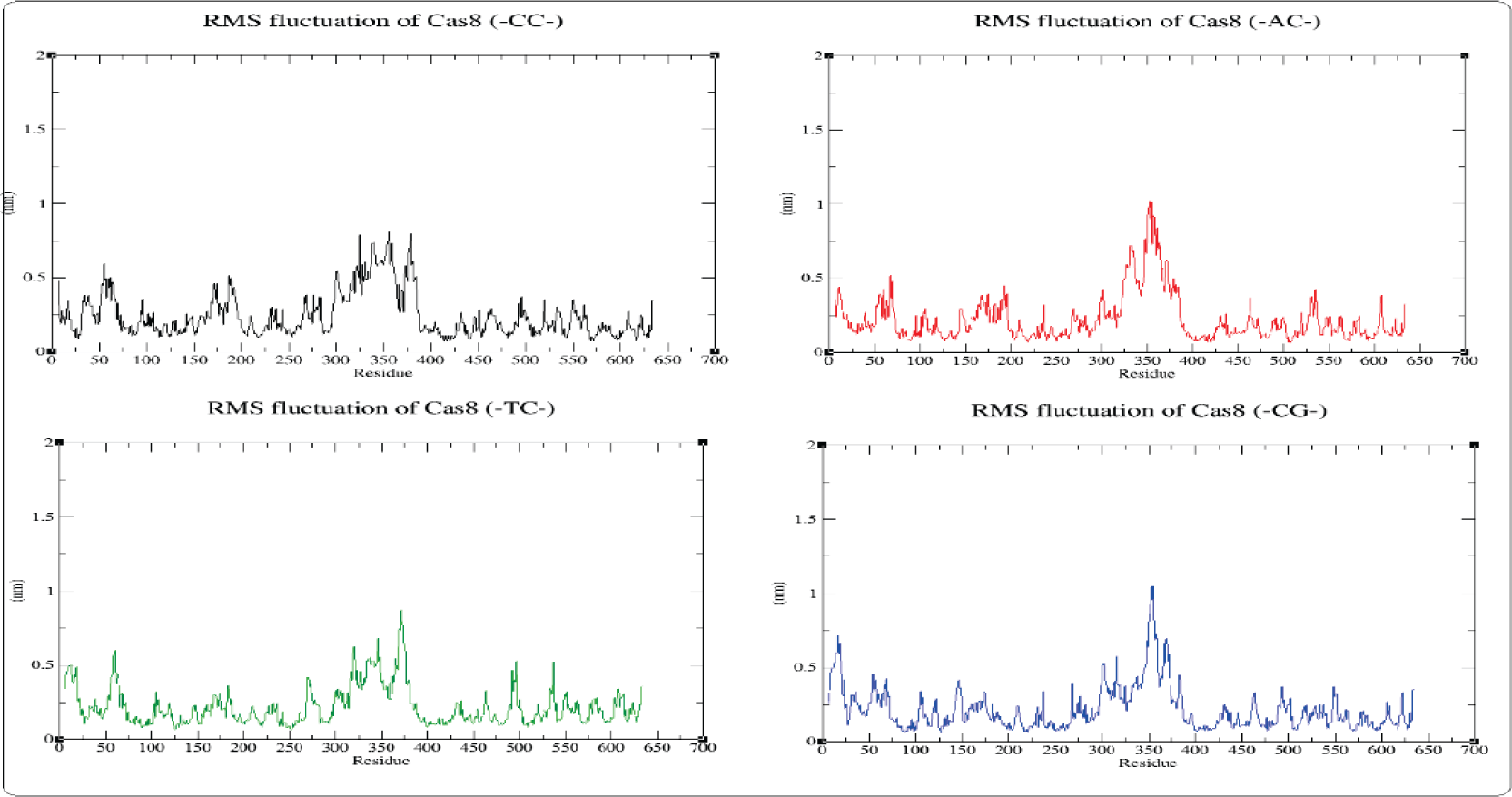
**Root means square fluctuation (RMSF) of Cas8 for each PAM model over the timescale of 1 μs simulation.**

The higher flexibility for all models was observed in the helical bundle regions located at aa 325–375, which is expected due to the intrinsic dynamic behavior of the Cas8-HB. Close observation of the RMSF plot analysis revealed that the loop region positioned at aa 64-70 is among the most flexible regions in all systems. The average RMSF values for the NTD are 0.225, 0.199, 0.196, and 0.192 nm in 5′-CC-3′, 5′-CG-3′, 5′-TC-3′ and 5′-AC-3′ PAM models, respectively. **As listed in Table 2**, the MD simulations identified the specific NTD residues that exhibited higher flexibility, with the 5’-AC-3’ PAM model displaying the fewest number of such residues. This suggests that the NTD’s flexibility is reduced upon binding to the 5’-AC-3’ PAM mutant, compared to the other model. Interestingly, the residues of the wild-type-PAM showed more dynamics and flexibility compared to other PAM models.

**Table 2.**
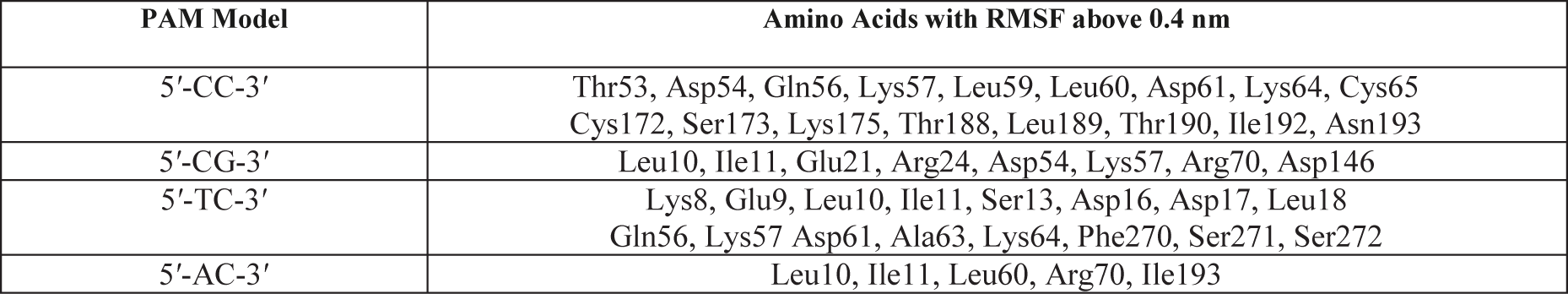
The N-terminal domain (NTD of Cas8 residues with RMSF values above 0.4 nm.

| PAM Model | Amino Acids with RMSF above 0.4 nm |
| --- | --- |
| 5'-CC-3' | Thr53, Asp54, Gln56, Lys57, Leu59, Leu60, Asp61, Lys64, Cys65<br>Cys172, Ser173, Lys175, Thr188, Leu189, Thr190, Ile192, Asn193 |
| 5'-CG-3' | Leu10, Ile11, Glu21, Arg24, Asp54, Lys57, Arg70, Asp146 |
| 5'-TC-3' | Lys8, Glu9, Leu10, Ile11, Ser13, Asp16, Asp17, Leu18<br>Gln56, Lys57 Asp61, Ala63, Lys64, Phe270, Ser271, Ser272 |
| 5'-AC-3' | Leu10, Ile11, Leu60, Arg70, Ile193 |

Hydrogen bonds play a critical role in stabilizing the secondary structures of proteins, including α-helices and β-sheets. They contribute to the protein’s rigidity and the selectivity of intermolecular interactions. In this study, the internal hydrogen bonds within the Cas8-PAM proximal region (first five nucleotides) were analyzed using the (*gmx hbond*) tool to determine the number of hydrogen bonds per time frame across all systems, **as shown in Fig. 10**

**Fig. 10:**
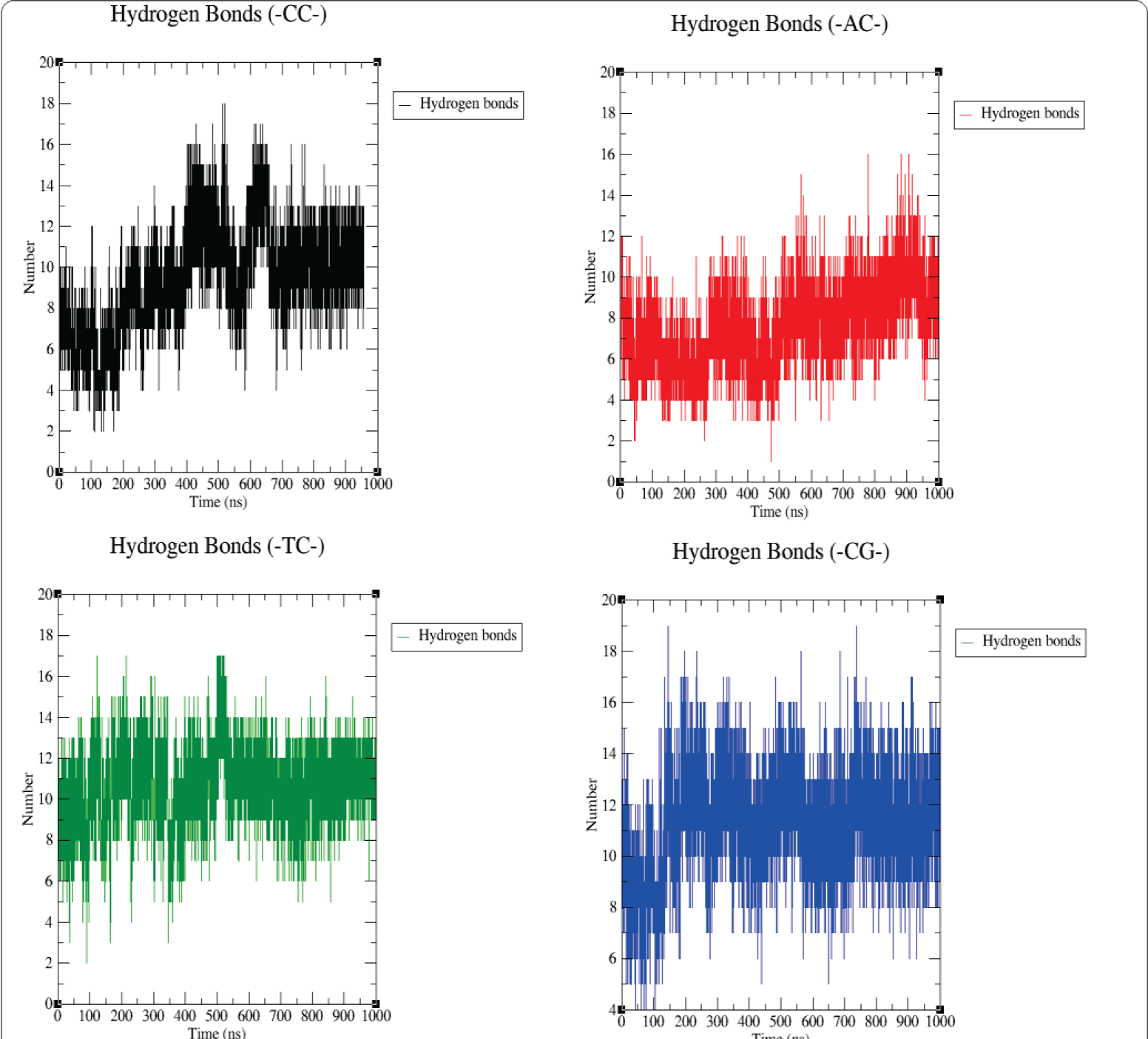
**The average number of hydrogen bonds within 0.35 nm between Cas8 and the PAM-proximal region over the timescale of 1 μs**

The average number of hydrogen bonds within 0.35 nm per frame between Cas8 and the PAM-proximal sequence for the 5′-CC-3′, 5′- CG -3′ and 5′-TC-3′ PAM models is 9.5, 11, and 10 respectively. The number slightly decreased to 7.5 per frame for the 5′-AC-3′ PAM model. More information about the hydrogen bonds and other nonbonded contacts has been discussed thoroughly in the subsequent sections.

### Interaction Analysis

To extract the most frequently visited conformations of Cas8 along the last 500 ns of each MD trajectory, an ensemble RMSD clustering algorithm (*gmx cluster*) was used to cluster the trajectories. This approach enabled the identification of representative structures that Cas8 most commonly adopted during the simulation. To examine the interactions between Cas8 protein and the PAM sequence, we employed the Gromos clustering algorithm using a cutoff of 0.3 nm. We then selected the top cluster, which accounted for 80% of the most commonly observed conformations, and analyzed it using the COCOMAPS (bioCOmplexes COntact MAPS) web application [23]. This tool highlighted the various interaction types between the protein and the PAM sequence within a distance of 0.8 nm, providing a comprehensive evaluation of the interactions in the system.

The minimum distances between any pair of atoms of the di-nt PAM sequence and Cas8 residues was computed. The analysis showed that the salt bridge is formed between the guanidinium side chain of the arginine (Arg246) and Guanine (DG53) in the case of 5′-CC-3′ and 5′-TC-3′ PAM models, and Cytosine (DC53) in the case of 5′-CG-3′ PAM model. The salt bridge involves interactions with the phosphate backbone through a combination of electrostatic attraction between a negatively charged phosphate group (labeled OP1 or OP2), and the positively charged guanidinium group (labeled as NH1 or NH2), and a hydrogen bond formed between the end group nitrogen in the guanidinium group (labeled as NH1 or NH2) and the side group oxygen on the phosphate (labeled O5 ’ or O4 ’), **as shown in Fig. 11-A, B, and C**. No electrostatic or hydrogen-bond interactions observed between Arg246 and DG53 in the 5′-AC-3′ PAM model, **as shown in Fig. 11-D**.

**Fig. 11.**
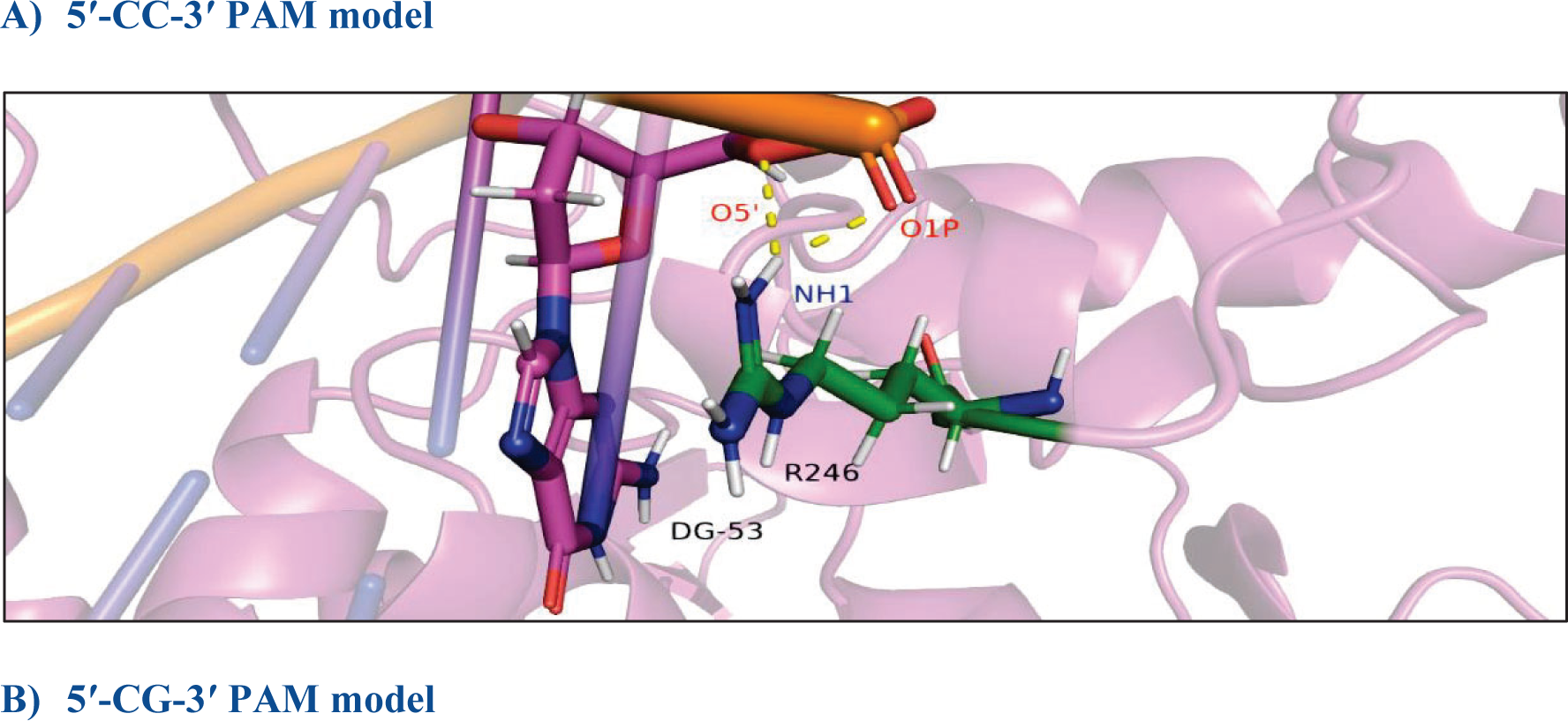
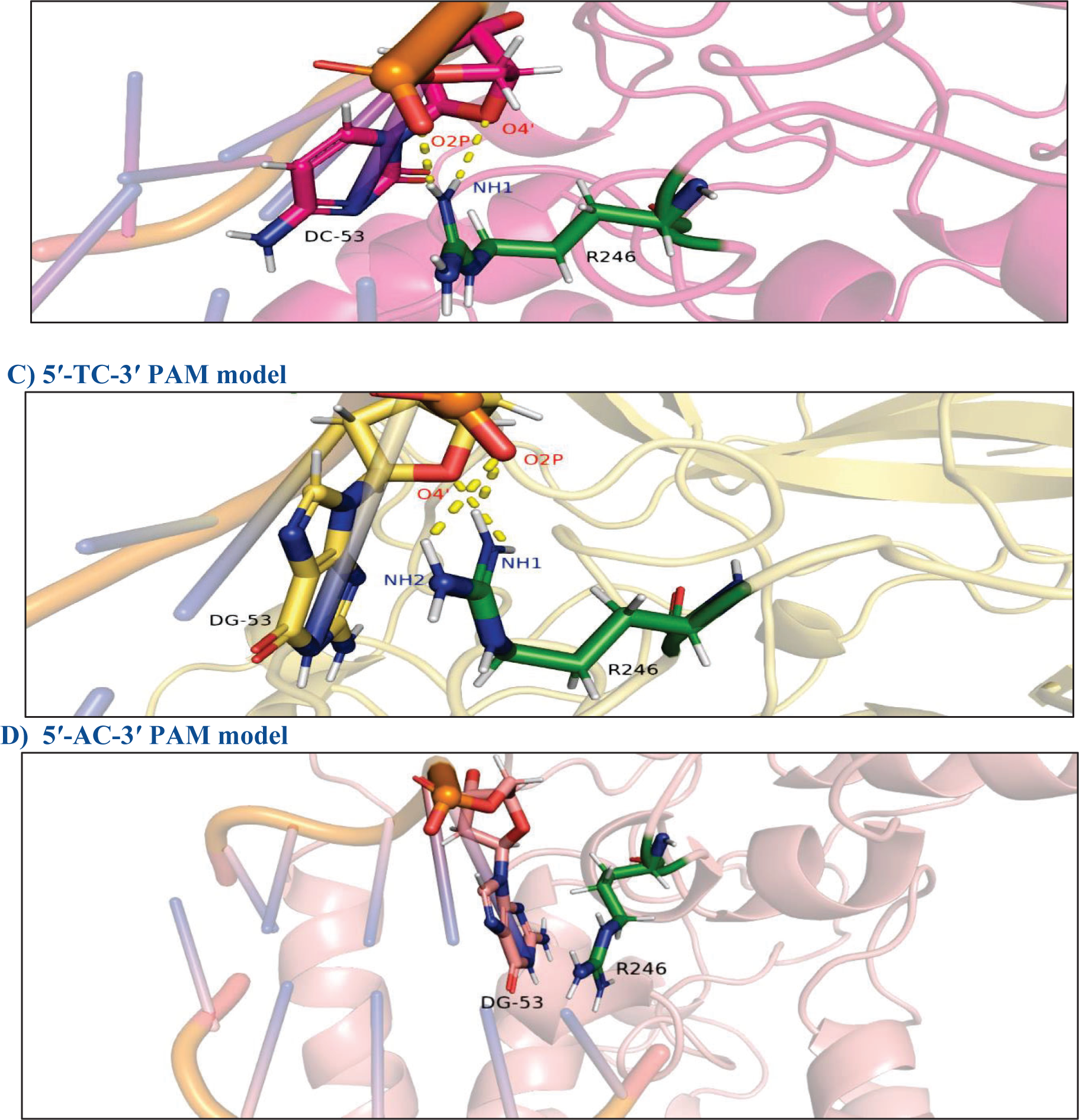
**The atomic interactions between the Arg246 and the PAM sequence.**

In the 5′-TC-3′ PAM model, a new hydrogen bond was observed between Alanine (Ala471, atom N) and Adenine (DA54, atom P). Similarly, in the 5′-AC-3′ PAM model, a new hydrogen bond was observed between Lysine (K462, atom NZ) and Guanine (DG53, atom O1P).

In the 5′-CC-3′ PAM model, Serine (Ser130) interacted with the DNA by forming a hydrogen bond with Cytosine (DC4). However, in the 5′-CG-3′ and 5′-TC-3′ PAM models, Serine 130 formed a hydrogen bond with Guanine (DG5) and Cytosine (DC5), respectively. No hydrogen bond was observed between Ser130 and the DNA in the 5′-AC-3′ PAM model, **as shown in Fig. 12**.

**Fig. 12.**
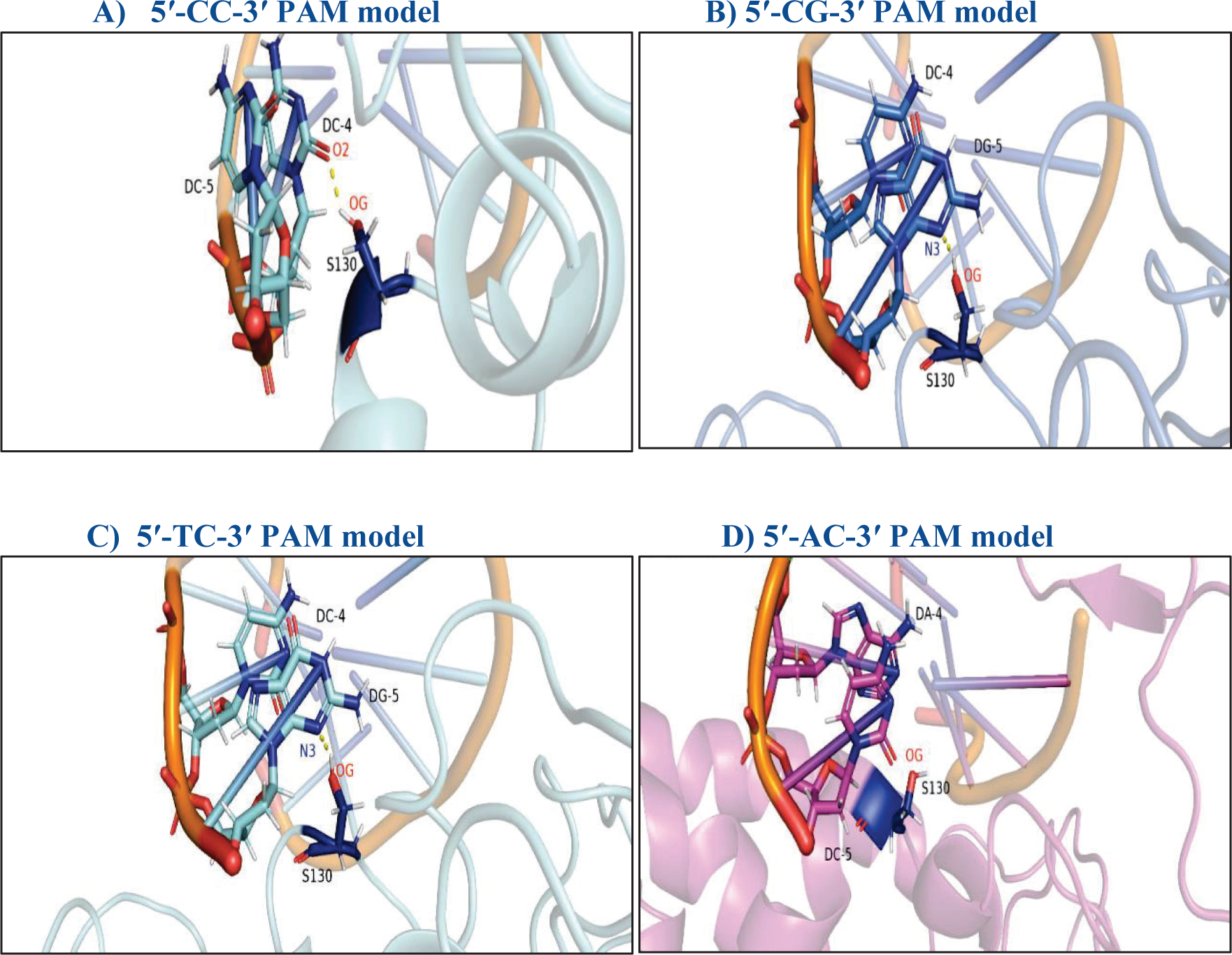
**The atomic interactions between the Ser130 and the PAM sequence.**

#### Minimum Distance Profile between Cas 8 Residues and the di-nt PAM Sequence

To explore the effects of PAM mutations on the distances between Cas8 residues and the di-nt PAM sequence, the minimum distance between Cas8 residues and the di-nt PAM was measured over time to assess changes in the dynamics of these residues during the simulation. The residues selected for analysis were Ser130, Arg246, and Lys462, based on the interaction analysis, **as shown in Fig. 13**

**Fig. 13:**
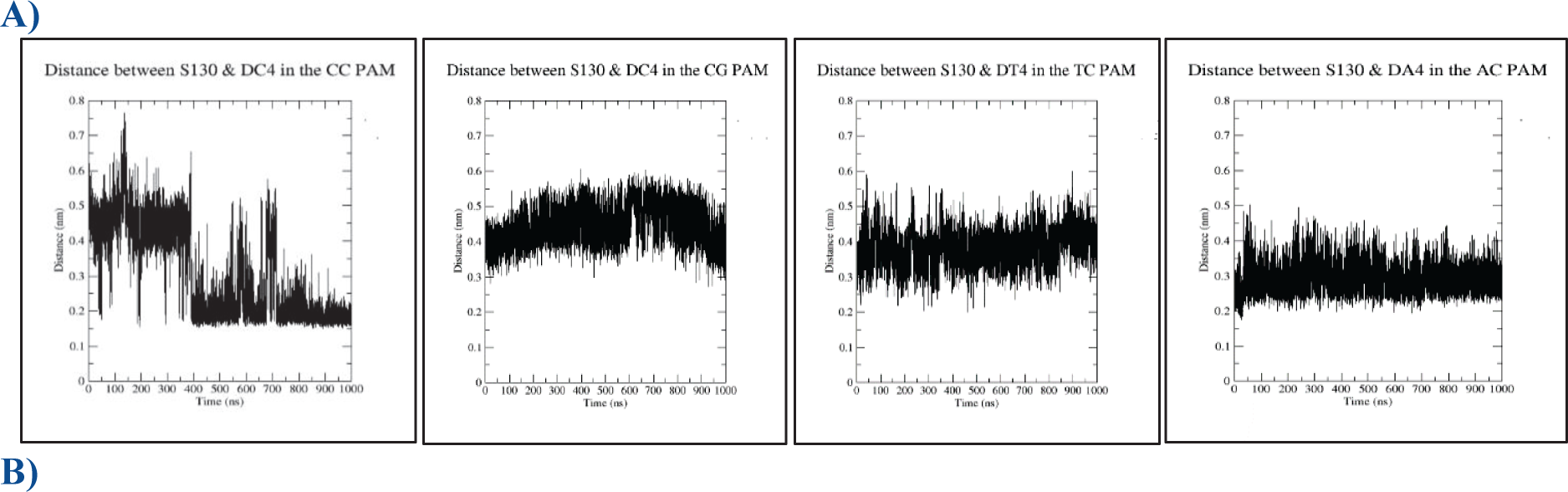
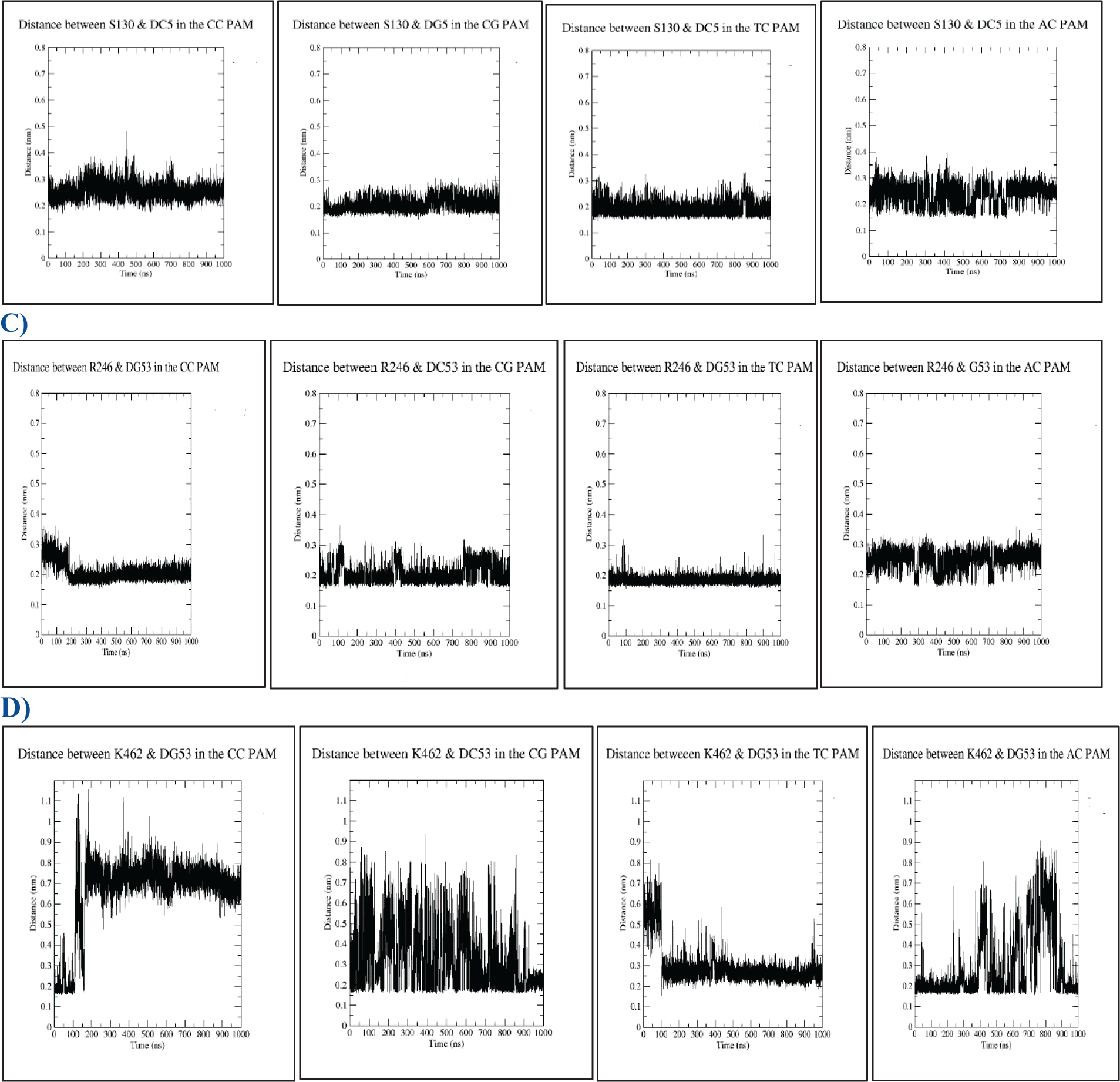
**The time evolution of the distance between Cas8 and the di-nt PAM sequence**

In the wild-type-PAM model, the distance between Ser130 and nt-4 varied greatly from 0 ns to 700 ns, ranging from 0.2 to 0.75 nm, before gradually decreasing to approximately 0.35 nm at around 750 ns. In all mutant models, the distance between Ser130 and nt-4 ranged from 0.2 to 0.6 nm throughout the entire simulation. Ser130 maintains contact with nt-5 with an average distance of ∼ 0.35 nm in the 5′-CC-3′, 5′-CG-3 and 5′-TC-3′ PAM models; although the distance profile indicated a tighter interaction for the 5′-CC-3′ and 5′-CG-3′ PAM models compared to the 5′-TC-3′ PAM model. Starting from 600 ns, the distance profile for the 5′-AC-3 PAM model revealed a very loose interaction between Se130 and nt-5.

For the 5′-CC-3′ PAM, the distance between the Arg246 and nt-53 was maintained within a range of 0.35 nm at the beginning of the simulation but gradually reduced to 0.16 - 0.26 nm starting from 200 ns simulation till the end of the simulation. A similar trend was observed for the 5′-CG-3′ and 5′-TC-3′ PAM models, in which the distance maintained within the cutoff of hydrogen bonds (0.35nm) throughout the simulation; however, the 5′-CG-3′ PAM model showed looser interaction. For the 5′-AC-3′ PAM model, although the distance was maintained within 0.35 nm, the interaction did not meet the hydrogen bond criteria.

In the 5′-TC-3′ PAM model, Lys462 interacts with DG53 with an average distance of 0.2-0.35 nm throughout the simulation. The interaction between Lys462 and nt-53 was tighter in the 5′-TC-3′ model compared to all other PAM models.

A difference contact analysis was conducted to determine how the mutations affected the distance between the atoms. The distances were computed based on the top clusters that represented nearly 80% of the most frequently visited conformations during the simulation. The computation of distances was performed using a cutoff of 0.8 nm. Residues that showed a significant decrease in distance indicated the formation of new interactions, while those that showed an increase in distance suggested a loss of interaction. **Tables S1-4** provide a complete list of the distances that were computed, whereas **Table 3** focuses on the most significant changes.

**Table 3.** The impact of PAM mutation was evaluated by calculating the difference in contact (in nm) between each mutant PAM model and the wild-type 5’-CC-3’ PAM model. This was achieved by subtracting the minimum distances in wild-type model from those in the PAM mutants models. The interactions that were significantly affected by the PAM mutation are presented. The dark red color indicates a newly formed interactions, and the dark blue color indicates a loss of interactions.

**A) Differences in distances between nt-54 in mutant models minus nt-54 in the 5'-CC-3' PAM**
|  |  | CG-CC | TC-CC | AC-CC |
| --- | --- | --- | --- | --- |
| LYS91 | nt-54 | 0.06 | -0.50 | -0.40 |
| ILE95 | nt-54 | -0.03 | -0.40 | -0.37 |
| ARG96 | nt-54 | -0.60 | 0.00 | 0.00 |
| HIS128 | nt-54 | 0.04 | -0.19 | -0.22 |
| LYS131 | nt-54 | -0.31 | -0.23 | -0.12 |
| TYR132 | nt-54 | -0.20 | 0.00 | 0.00 |
| SER245 | nt-54 | -0.28 | 0.00 | 0.00 |
| ARG246 | nt-54 | 0.00 | 0.26 | 0.03 |
| THR247 | nt-54 | -0.01 | -0.14 | 0.02 |
| THR248 | nt-54 | 0.27 | 0.34 | 0.35 |
| MET250 | nt-54 | 0.10 | 0.26 | 0.08 |
| LYS462 | nt-54 | 0.33 | -0.29 | -0.21 |
| SER468 | nt-54 | 0.55 | 0.33 | 0.49 |
| ALA469 | nt-54 | 0.43 | 0.20 | 0.58 |
| PRO470 | nt-54 | 0.04 | -0.25 | 0.11 |
| ALA471 | nt-54 | -0.10 | -0.22 | 0.19 |

**B) Differences in distances between nt-4 in mutant models minus nt-4 in the 5'-CC-3' PAM**
|  |  | CG-CC | TC-CC | AC-CC |
| --- | --- | --- | --- | --- |
| LYS52 | nt-4 | -0.29 | 0.00 | 0.00 |
| GLN67 | nt-4 | -0.50 | 0.00 | 0.00 |
| LYS68 | nt-4 | -0.52 | 0.00 | 0.00 |
| GLU72 | nt-4 | -0.16 | 0.00 | 0.00 |
| ASP129 | nt-4 | 0.18 | 0.09 | 0.01 |
| SER130 | nt-4 | 0.29 | 0.24 | 0.13 |
| ASN134 | nt-4 | 0.22 | 0.27 | -0.04 |
| LYS135 | nt-4 | 0.20 | -0.04 | 0.17 |
| THR248 | nt-4 | 0.22 | 0.30 | 0.16 |

**C) Differences in distances between nt-53 in mutant models minus nt-53 in the 5'-CC-3' PAM**
|  |  | CG-CC | TC-CC | AC-CC |
| --- | --- | --- | --- | --- |
| ILE95 | nt-53 | 0.00 | -0.41 | -0.31 |
| ARG96 | nt-53 | 0.00 | -0.32 | -0.39 |
| SER130 | nt-53 | -0.03 | -0.30 | -0.37 |
| LYS131 | nt-53 | 0.00 | -0.15 | -0.21 |
| SER245 | nt-53 | -0.30 | -0.12 | -0.02 |
| THR247 | nt-53 | 0.14 | 0.08 | 0.15 |
| THR248 | nt-53 | 0.26 | 0.04 | 0.03 |
| MET250 | nt-53 | 0.19 | 0.05 | 0.04 |
| PHE459 | nt-53 | 0.00 | -0.14 | 0.00 |
| VAL460 | nt-53 | 0.00 | -0.21 | -0.26 |
| LYS462 | nt-53 | 0.16 | -0.46 | -0.55 |
| SER468 | nt-53 | -0.03 | 0.30 | -0.04 |
| ALA469 | nt-53 | 0.30 | 0.53 | 0.16 |
| PRO470 | nt-53 | -0.09 | -0.35 | -0.35 |
| ALA471 | nt-53 | -0.29 | -0.34 | -0.18 |
| THR472 | nt-53 | -0.34 | -0.37 | -0.14 |

**D) Differences in distances between nt-5 in mutant models minus nt-5 in the 5'-CC-3' PAM**
|  |  | CG-CC | TC-CC | AC-CC |
| --- | --- | --- | --- | --- |
| LYS23 | nt-5 | 0.39 | 0.20 | -0.01 |
| ARG27 | nt-5 | 0.26 | 0.18 | 0.07 |
| LEU49 | nt-5 | 0.19 | 0.19 | -0.01 |
| THR50 | nt-5 | 0.00 | -0.23 | 0.00 |
| ASP51 | nt-5 | 0.00 | -0.23 | -0.03 |
| THR53 | nt-5 | 0.00 | 0.00 | -0.16 |
| SER130 | nt-5 | -0.03 | -0.09 | 0.54 |
| VAL133 | nt-5 | -0.20 | 0.00 | -0.23 |
| LYS135 | nt-5 | 0.22 | -0.02 | 0.25 |
| ARG246 | nt-5 | -0.39 | -0.17 | -0.35 |
| ASN249 | nt-5 | -0.32 | -0.10 | -0.07 |
| MET255 | nt-5 | -0.25 | -0.17 | -0.29 |

The analysis of the contact map revealed that in the PAM models with 5’-TC-3’ and 5’-AC-3’ sequences, there was a noticeable reduction in the interaction distance between Lys91, His128, Ile95, and Lys462 residues with nt-54. On the other hand, Thr248, Ser468, and Ala469 residues showed the most elongation across all PAM models. In the case of Asn134, which interacts with nt-4, there was an increase in distance in the 5’-CG-3’ and 5’-TC-3’ models, but a slight decrease in the 5’-AC-3’ PAM model. Furthermore, in the 5’-TC-3’ and 5’-AC-3’ PAM models, the distance between Ile95, Arg96, Ser130, Val460, Lys462, and Pro470 residues that interact with nt-53 decreased upon mutation. Finally, there was a shortening of the distance between Thr53 and nt-5 in the 5’-AC-3’ PAM mutation, while Ser130 showed the greatest elongation.

The analysis indicated that the distance between the di-nt PAM sequence and Cas8 residues exhibited significant variation due to the mutation. Specifically, the distance between Val460, Lys462, and Pro470 decreased as a result of a mutation in the 5′-TC-3′ and 5′-AC-3′ PAM, indicating the formation of new interactions. It is worth noting that these two PAM mutants did not achieve complete integration, suggesting that these new interactions may have a negative impact on integration efficiency.

### Effect of PAM Mutation on the Allosteric Communication between Cas8 Residues

PAM binding can cause changes in the conformation of Cas8, which may propagate through the protein along allosteric pathways, triggering a cascade of events to activate the transposition process. Different analysis approaches were used to identify the residues involved in the allosteric pathway upon PAM binding. The first approach used the perturbation response scanning method (PRS) using the DynOmics ENM web server [24]. The second approach used the dynamics residues network (DRN) to compute relevant information using the MDM-TASK [25] . The third approach computed the correlation of Kapsch-Sander (KS) electrostatic energy throughout the MD simulation trajectories using a novel method known as MDiGest [26].

The DynOmics utilizes a network model consisting of nodes representing the amino acid residues and edges defining the strength of correlations established between the residues [24,27] . The top cluster of each PAM model was uploaded into the DynOmics elastic network model (ENM) web to investigate the capability of residues to act as potential mediators of allosteric communication based on the ability to work as “effector residues “ that respond to the external perturbations and propagate the signals to the “sensor residues” that sense the motion and alter the dynamics [28] .

The evaluation of residues is conducted using the perturbation response scanning method (PRS), which enables a quantitative assessment of the perturbation response of each residue to every other residue. PRS measures the response strength of the “sensor” residue i to perturbations in the “effector” residue j [28]. Sensors and effectors residues in each system **are shown in Fig. 14**

**Fig. 14.**
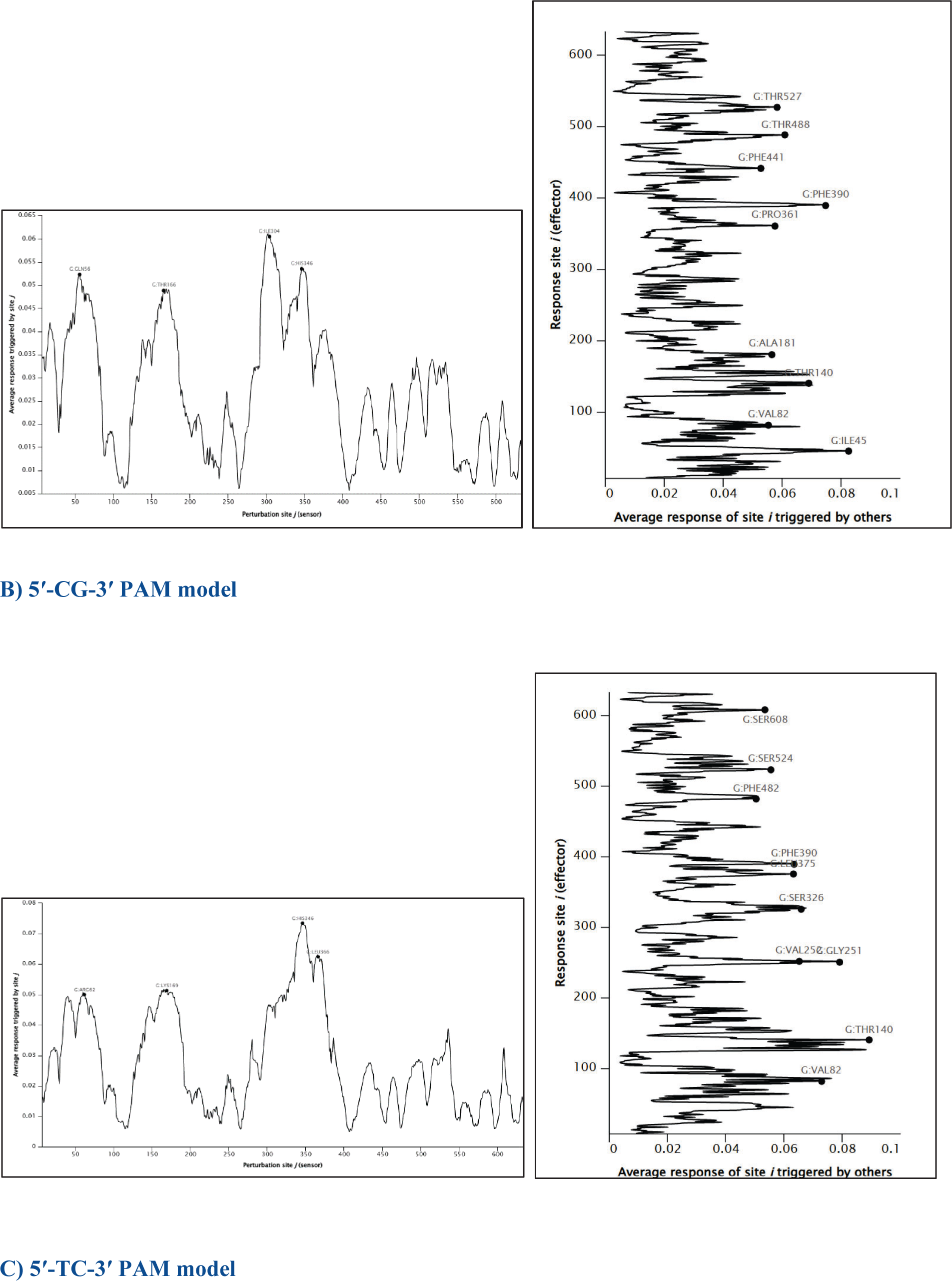
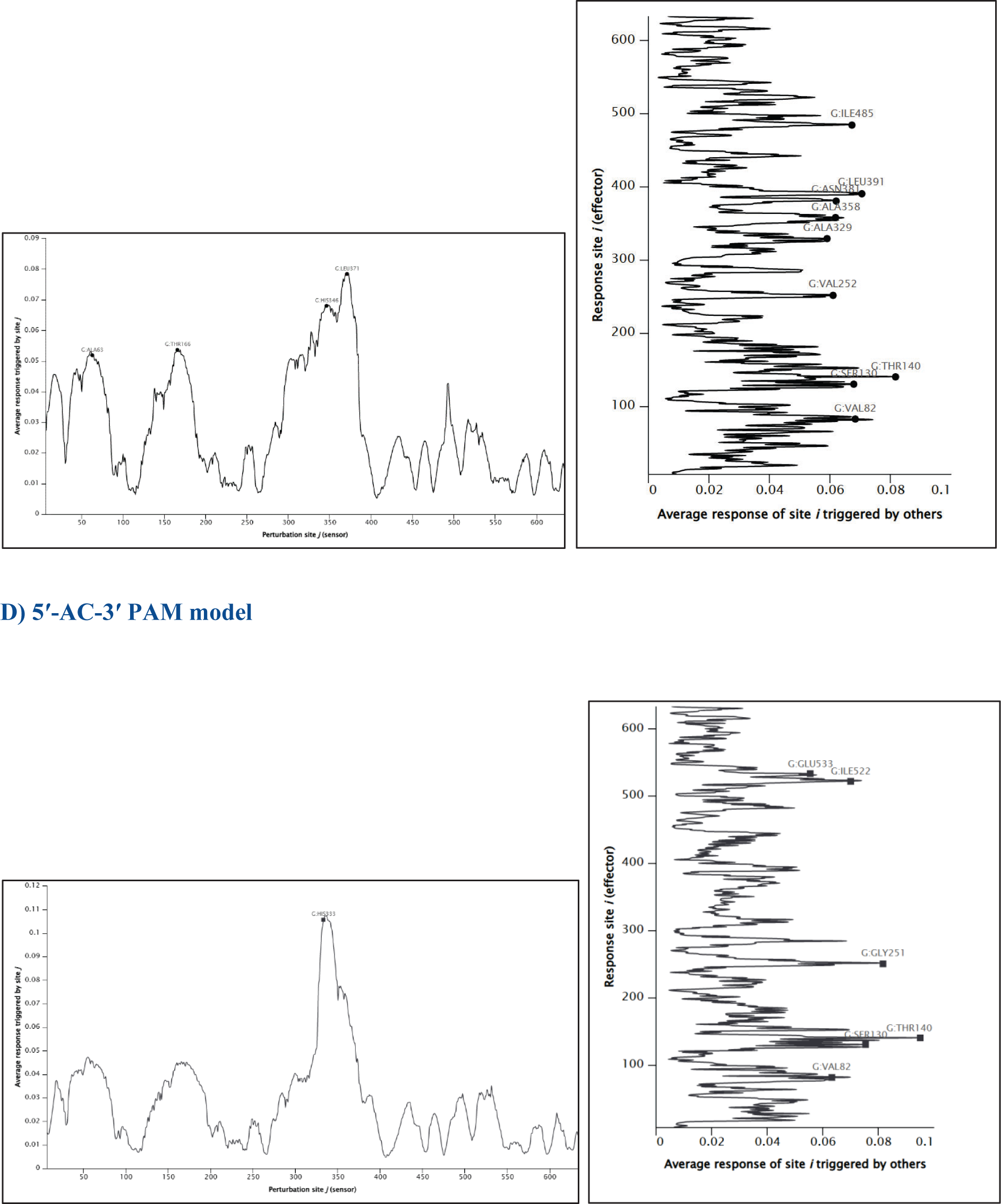
**The propensity of the Cas8 residues to act as sensors or effectors in the wild-type-PAM and mutant PAM models.**

**Fig. 15.**
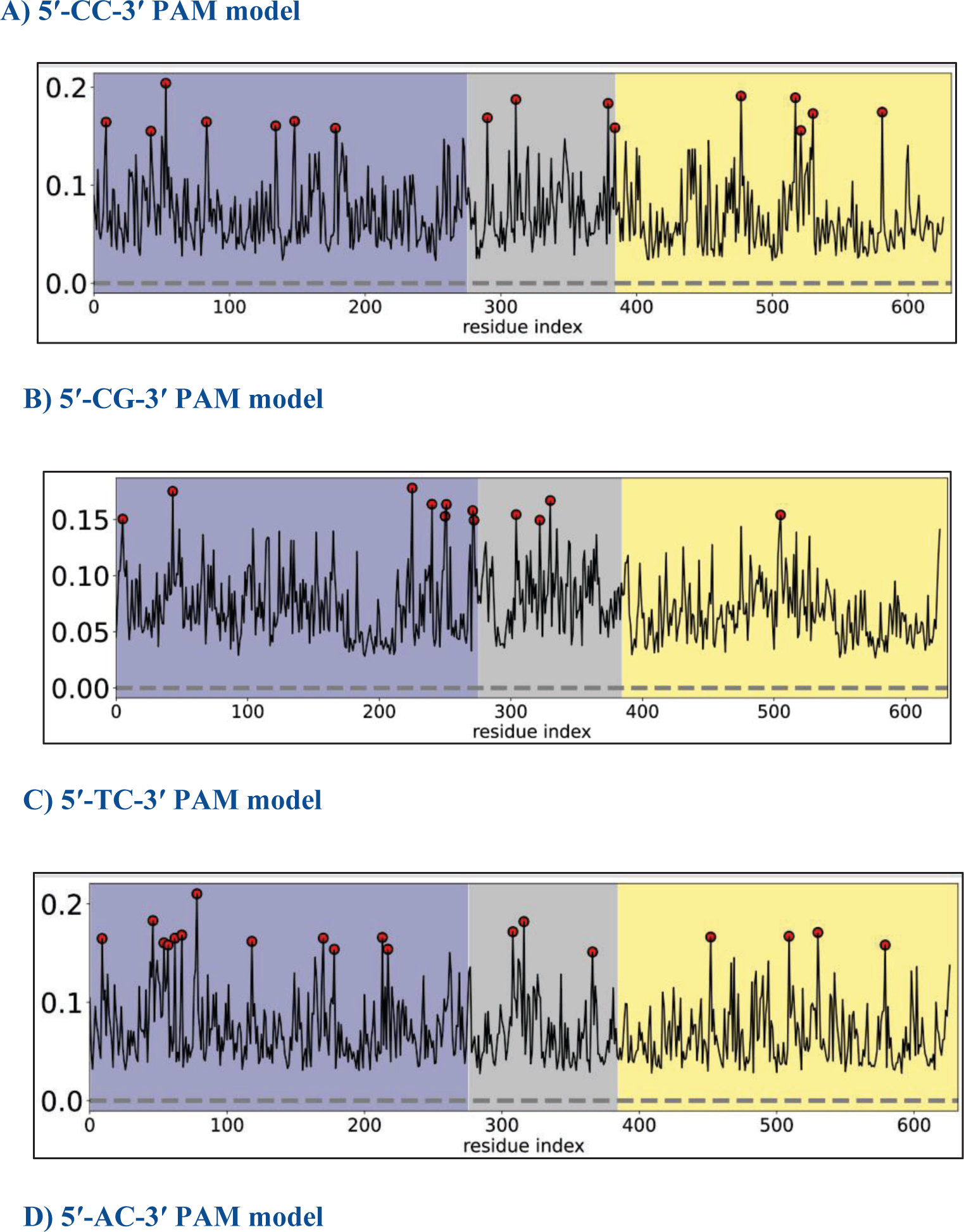
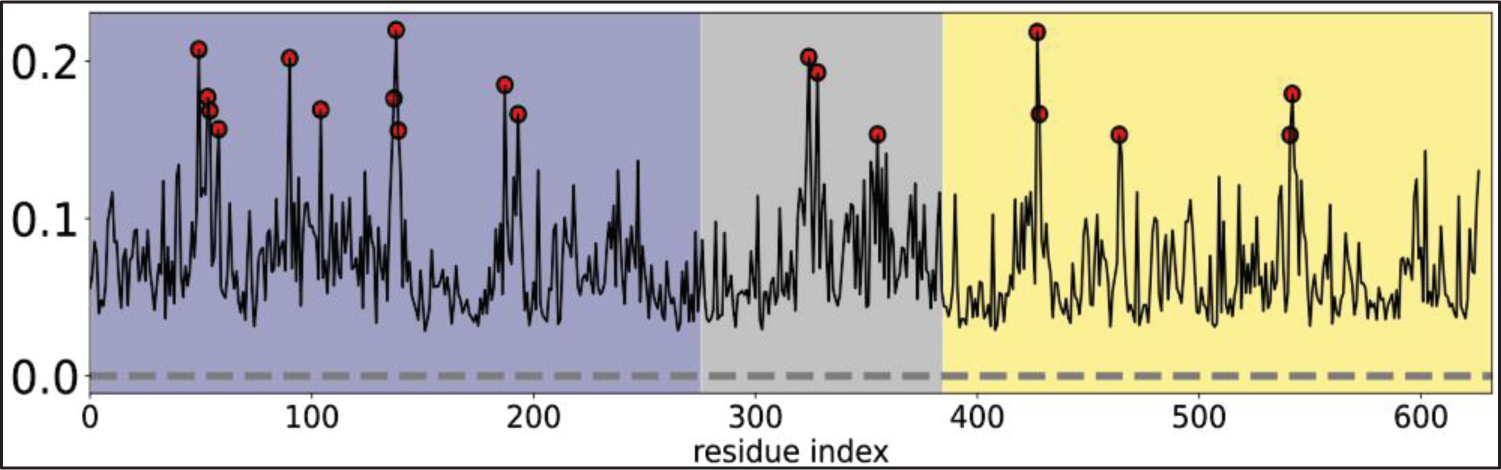
**Cas8 residues with high eigen centrality values upon binding the PAM sequence.**

Effector residues are believed to participate in the PAM binding process and can effectively transmit signals to control downstream effects [24]. In contrast, sensor residues, which are located at distant sites, are sensitive to allosteric perturbations and are associated with changes in local conformation as a response to such perturbations [24]. Peaks in the average profile indicate sensor and effector residues that are anticipated to undergo significant displacements [24].

The analysis revealed that all of the PAM models, except for the 5’-AC-3’ model, had multiple sensor peaks that overlapped. These shared regions comprised a loop at position 56-66 and a helix at position 160-171. Additionally, the loop at position 340-354 was a common site for overlapping sensor residues across all PAM models and was determined to be the most sensitive region to perturbations. This region is a component of the Cas8-HB, which is believed to trigger TniQ activation. Furthermore, the investigation revealed that while certain effector residues, such as Val82 and Thr140, were present in all PAM models, each model had its own distinct set of effector residues that were unique to that model.

To compute the dynamic residue network (DRN), the second method involved extracting the final 500 ns of every simulation and analyzing them using MDM-TASK-web scripts that utilize a betweenness centrality metric [25]. Betweenness centrality (BC) is a metric used to assess the flow of information in a graph through a network model, focusing on how often a node acts as a bridge along the shortest path between two other nodes. It is based on the number of shortest paths that pass-through a given node. Residues with higher betweenness centrality would have greater control over the network, indicating a greater flow of information through those residues [25]. The following residues have high average BC values upon binding to all PAM sequences: Arg85, Gln86, His88, Val398, His423 and Gly572, Glu595, Trp617, and Val628, indicating that they play a critical role in controlling the flow of information within the network. This finding suggests that these residues are likely to be involved in the allosteric regulation of Cas8.

A third method was employed to conduct further analysis to calculate the correlation throughout a molecular dynamic’s trajectory. This method utilized the Kapsch-Sander (KS) electrostatic energy and was performed using a new approach called MDiGest [26,29]. In order to evaluate the electrostatic energy between pairs of Cas8 residues upon binding to each PAM sequence, KS electrostatic energies were computed between every pair of residues in each frame. This was done by quantifying the electrostatic energy between the acceptor carbonyl (CO) backbone group and the donor amine (NH) backbone groups. The resulting correlation coefficient was used to generate a matrix that represented the electrostatic coupling between each donor/acceptor pair of Cas8.

Next, the eigenvector centrality (EC) metric was used to identify key residues that played a significant role in the propagation of electrostatic information throughout Cas8. This metric relied on the fact that central residues with more neighbor residues were highly connected and influential in the graph [26,30]. The residues with high eigen centrality values are circled in red, **as shown in**

Residues in Cas8 with high eigen centrality were found to be similar across all PAM models, indicating that PAM mutations did not have a significant impact on the propagation of information within Cas8. However, His88, Val398, Gly572, and Glu595 were consistently identified as key residues in various analyses.

The MDiGest tool uses community network analysis to group residues based on their shared secondary structure during an MD simulation. To allow for comparison between the Cas8 protein when it is unbound and when it is bound to each PAM sequence, an additional MD simulation of an apo system without DNA (PDB Code: 6V9Q) was conducted [7]. The edges between the communities in the community network represent the correlation coefficients calculated from the previously KS energies obtained, **as shown in Fig. 16**

**Fig. 16.**
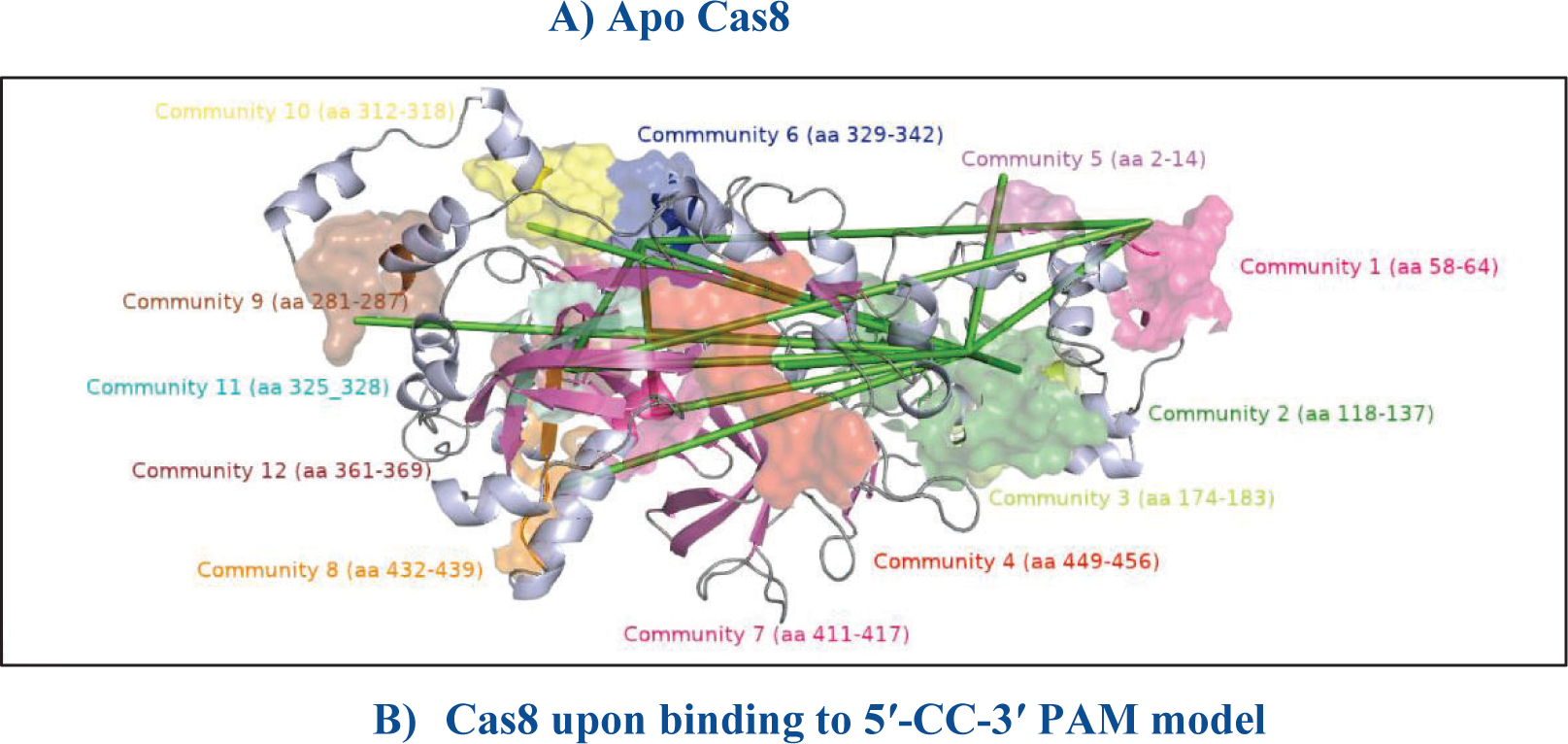
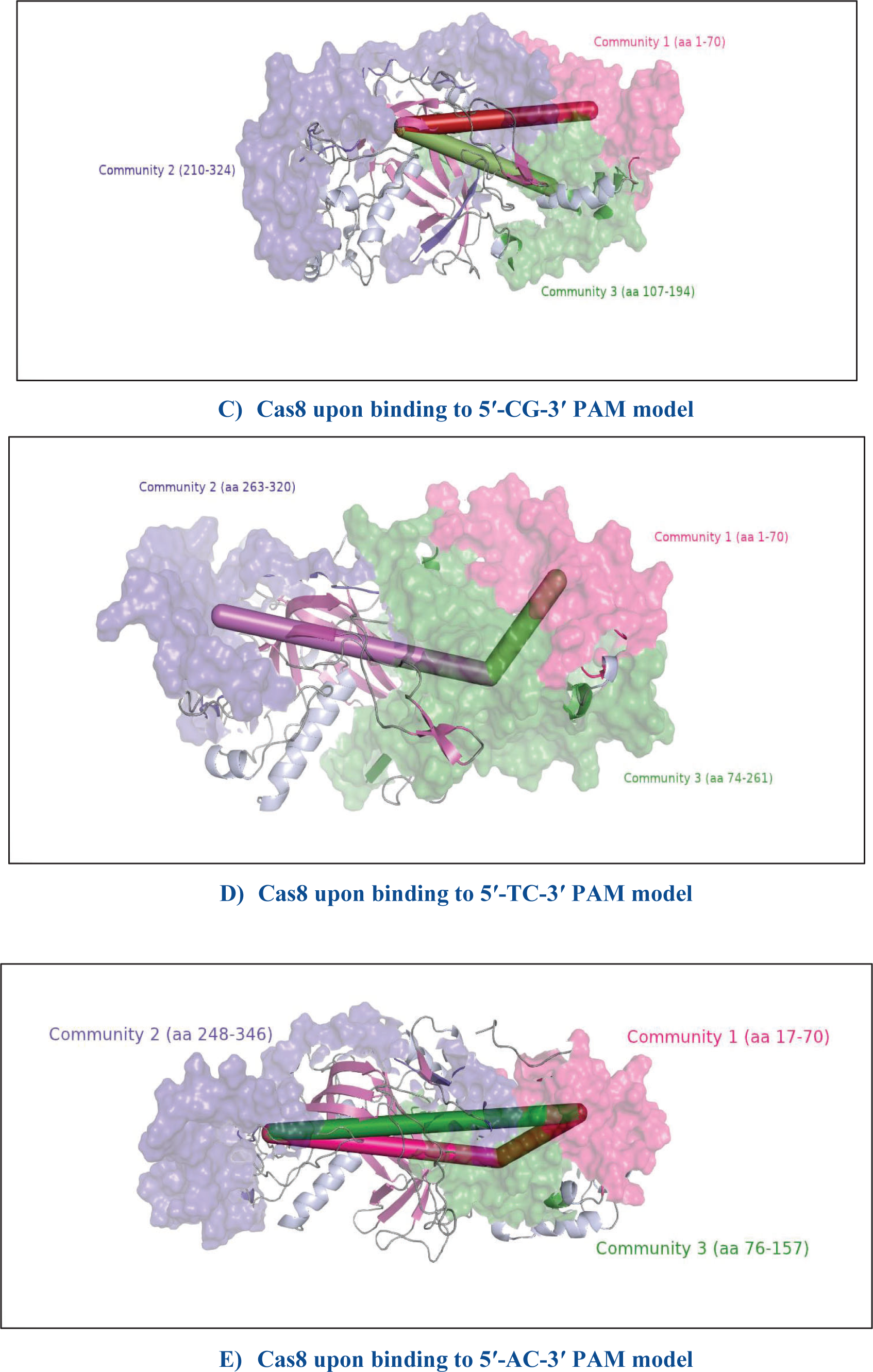
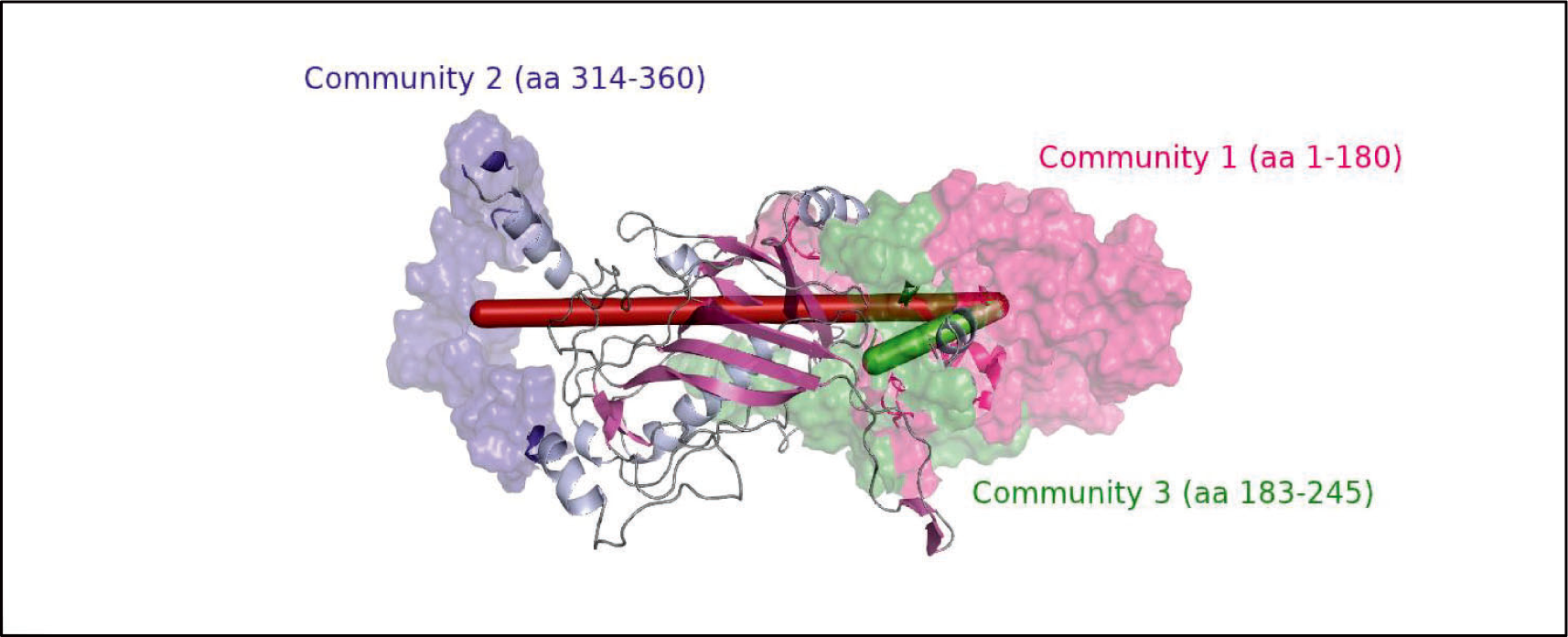
Community network analysis of Cas8 in the apo system and the PAM-bound Cas8. Created with BioRender.com

The analysis showed that the apo Cas8 had 13 fragmented communities with a weak correlation between Cas8 residues. However, upon PAM binding, the correlation between residues was strengthened, and the number of communities decreased to three, regardless of the PAM sequence.

The analysis further revealed that PAM induces a more robust allosteric communication between the amino acid residues belonging to the N-terminal domain (community 1) and the helical bundle domain of Cas8 (community 2). The community networks provide a clear indication of which regions of Cas8 are strongly coupled and how allosteric communication is affected by PAM binding.

## Discussion

According to the results, the RMSD and RMSF plots of all mutant models were similar to those of the wild-type PAM model, which suggests that the stability and flexibility of Cas8 were not influenced by the mutations. Both 5′-CC-3′ and 5′-CG-3′ PAMs demonstrated high integration efficiency upon binding to Cas8. However, the 5′-CG-3′ PAM was found to have greater conformational variability, indicating lower stability and specificity compared to the wild-type PAM.

The analysis of Cas8 interactions showed that several critical residues (Ser130, Arg246, Thr248, Asn249) are involved in PAM recognition. In type I CRISPR systems, Cas8 scans the DNA to identify the correct PAM sequence, and Arg246 serves as a wedge to separate the double-stranded DNA and displace the non-target strand from the target strand. The Arg246 wedge causes the first protospacer nucleotide in the target strand to rotate outward. However, in the 5’-AC-3’ PAM model, the Arg246 orientation is different, and this may prevent Arg246 from acting as a wedge and separating the double-stranded DNA, **as shown in Fig. 17**

**Fig. 17.**
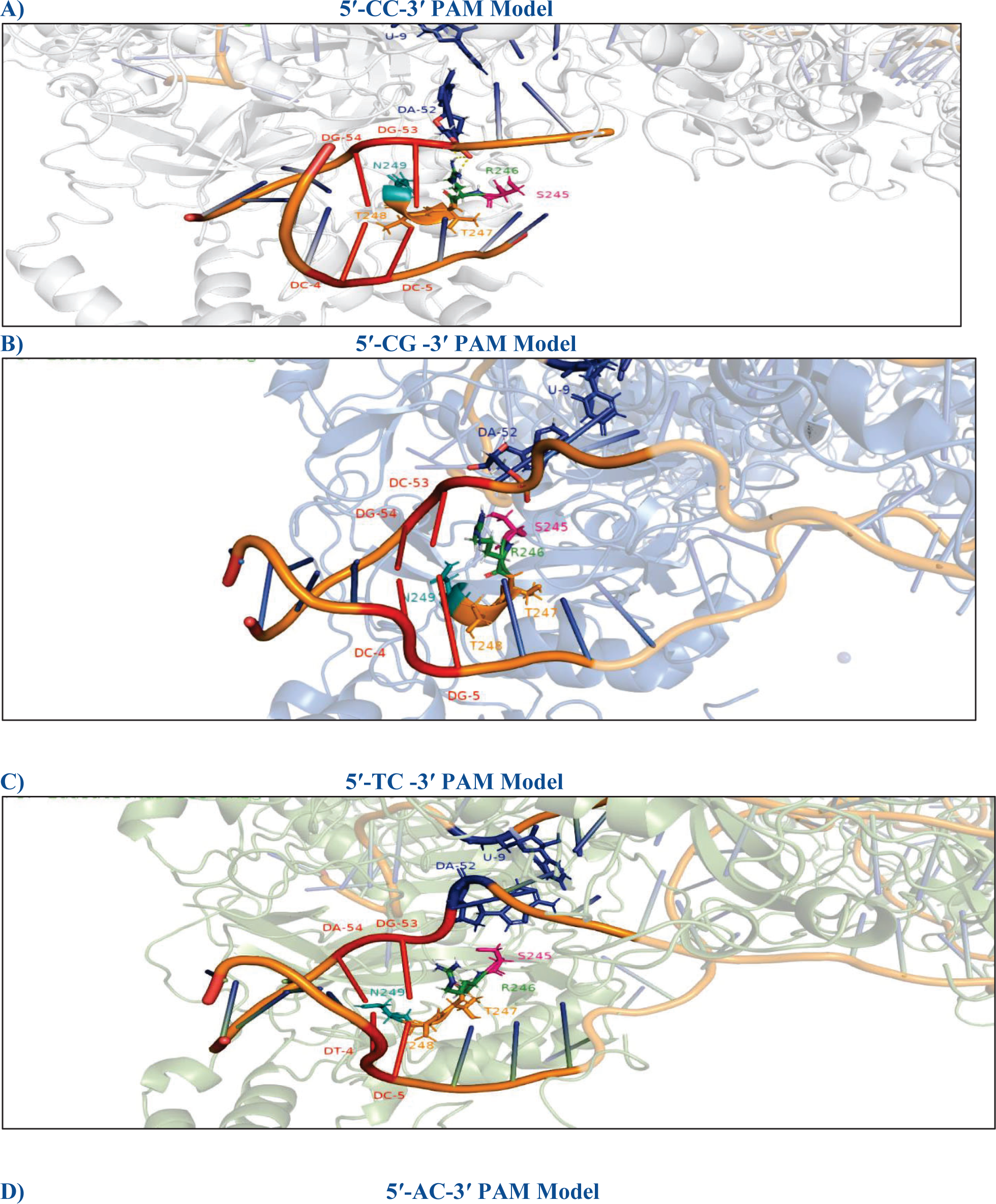
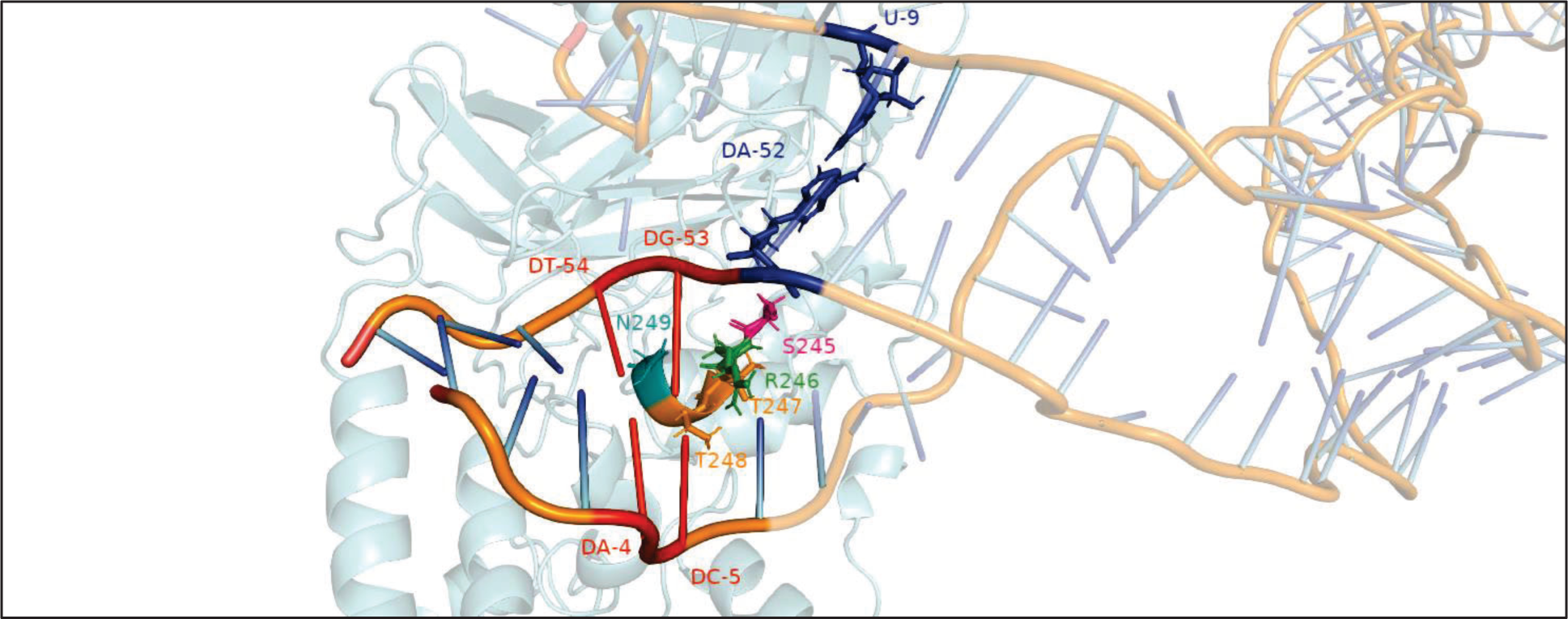
**Arginine wedge upon binding to each PAM sequence**

The salt bridge formed between the critical arginine residue Arg246 in the binding site of the N-terminal domain (NTD) of Cas8 and the PAM sequence is an important interaction involved in PAM recognition. The 5’-AC-3’ PAM mutation breaks this key contact, which is likely the primary reason for the lack of integration. The salt bridge interaction between Arg246 and the PAM sequence (specifically DG53) helps to position the Cas8 protein correctly for PAM recognition and to initiate the DNA unwinding process. Without this crucial interaction, the Cas8 protein may not be able to correctly recognize or interact with the PAM sequence, which could lead to a decrease in integration efficiency or even complete inhibition.

**As shown in Fig. 12**, another important interaction to consider is the interaction between Ser130 and DNA. Upon inspection, it was observed that the side chain of Ser130 mainly interacts with the Cytosine (DC4) base in the wild-type 5′-CC-3 PAM model throughout most of the simulation time. However, in the 5′-CG-3′ and 5′-TC-3′ PAM models, Ser130 interacts with nt-5 instead. In the 5′- CG-3′ PAM model, the distance analysis revealed that Ser130 is closer to Guanine (DG5) within the cutoff distance for hydrogen bonds. This same scenario was observed in the 5′-TC-3′ PAM model, where Ser130 interacts with the Cytosine (DC5) and forms a hydrogen bond for 87% of the simulation time. In contrast, in the 5′-AC-3′ PAM model, the side chain of Ser130 is rotated outward, which prevents any interaction with either nt-4 or nt-5.

The distances between Val460, Lys462, and Pro470, and the PAM region showed a significant reduction in the 5′-TC-3′ and 5′-AC-3′ PAM mutants. We believe that this reduction had a negative impact on the integration process as it caused the key residues to move away from the PAM region, disrupting the interaction and ultimately leading to unsuccessful integration.

In the allosteric analyses conducted using the three approaches mentioned, it was found some amino acids - Val82, His88, Thr140, and Phe390, Val398, His423, Gly572, and Glu595 - appeared in all four PAM models. These amino acids could be allosteric hotspots because of their influence on other residues. The fact that these amino acids are present in all PAM models, regardless of whether the PAM is recognized or not, as in the 5′-AC-3′ PAM, suggests that they play an important role in the overall function of the Cas8 protein.

The secondary structure community network analysis showed a weak coupling between communities, in the apo model, due to the fragmentation of communities. The number of communities is reduced upon PAM binding, even in cases where the PAM sequence is 5’-TC-3’ and 5’-AC-3’, with integration rates of 60% and 0%, respectively. This is because the PAM sequence strengthens the correlation between Cas8 residues, resulting in a more cohesive and strongly coupled system. As a result, the residues are grouped in fewer, more tightly correlated communities rather than being fragmented into multiple weakly correlated communities. This suggests that PAM binding induces a more efficient and coordinated allosteric communication among the Cas8 amino acid residues, which is important for the proper functioning of the CRISPR system. Our findings indicate that the PAM sequence serves two purposes in the INTEGRATE system. Firstly, it initiates the process by identifying the target DNA. Secondly, it acts as an allosteric regulator by facilitating communication between the N-terminal and Cas8-helical bundle domains.

## Conclusion

The collaboration between Tns proteins and Cas genes in transposon-encoded-CRISPR-Cas systems decreases the chance of introducing underinsured mutations by integrating the transposon at specific locations [5]. This upgraded CRISPR-Cas system could be very efficient as a gene editing tool. However, more work is needed to understand the molecular mechanisms of this system before applying it as a gene editing tool. This study focused on the PAM recognition step, the primary step in the integration process. We have comprehensively explored the mechanism of PAM recognition of the wild type 5′-CC-3′ PAM and three mutants (5′-CG-3′, 5′-TC-3′, and 5′- AC-3′ PAMs) through all-atoms MD simulations. The study identified the critical contact essential for the PAM recognition process, which is mainly mediated by Ser130 and Arg246. The simulations revealed that the interaction between these two residues was disrupted in the 5′-AC-3′ PAM mutant model. Notably, Arg246 adjusts its position to maintain contact with the DNA in all PAM models except the 5′-AC-3′ PAM, which could explain the observed failure of integration in this mutant. Our study investigated the role of the PAM sequence in the INTEGRATE system and its effect on the integration process. The results showed that the PAM sequence plays two crucial roles in the INTEGRATE system: initiating the integration process by recognizing the target DNA and acting as an allosteric regulator by facilitating communication between the N-terminal domain and the Cas8-helical bundle domain. The distance between the Cas8 residues and the PAM sequence is critical in ensuring successful integration, particularly in the presence of mutations. The study also identified some amino acids that could be classified as allosteric hotspots, because of their ability to significantly affect all other nearby residues. Although the significance of these particular residues has not been tested, they could be promising candidates for future mutagenesis studies. Overall, this study has provided valuable insights into the molecular mechanisms of the INTEGRATE system and the basis for allosteric communication within Cas8. Further research is needed to fully understand the mechanisms involved in the integration process and optimize the INTEGRATE system for practical applications.

## Acknowledgement

Our research was made possible through the use of computational resources provided by the Shaheen supercomputer at King Abdullah University of Science and Technology (KAUST) and the Bridges-2 supercomputer at the Pittsburgh Supercomputing Center. We express our gratitude for generously providing the essential computational resources required for our research.

## Ethics approval and consent to participate

Not applicable

## Consent for publication

Not applicable

## Availability of data and material

Not applicable

## Competing interests

No potential competing interest was reported.

## Funding

No funding was received to support the review article presented in the manuscript submitted.

## Authors’ contributions

All authors reviewed and approved the manuscript submitted.

## Supplementary

**Table S1:**
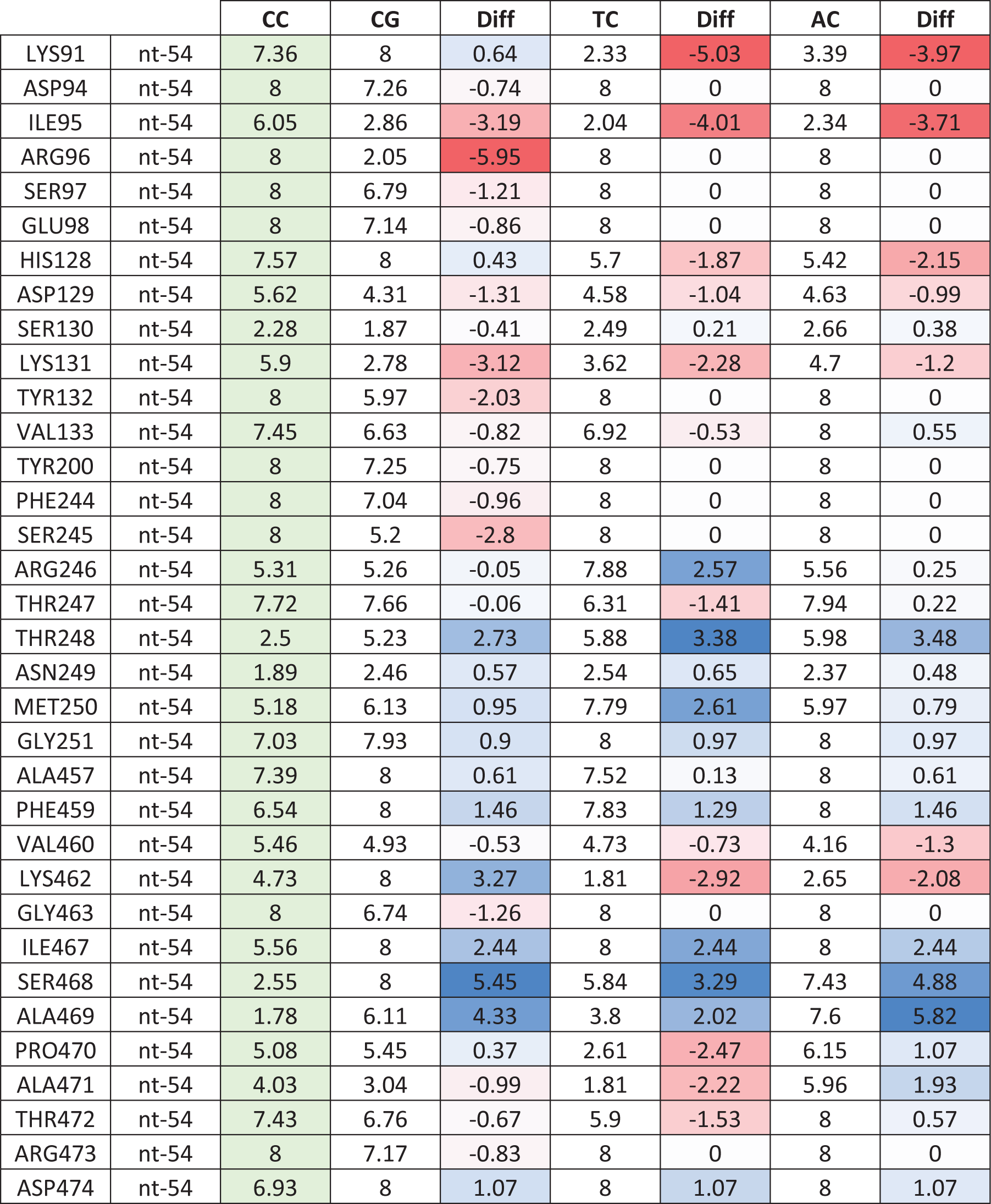
The Difference contact map between Cas8 and nt-54 in the PAM sequence in Angstrom (Å). Red color represents Negative values indicating a shortening in the distances upon mutation, whereas blue color represents positive values indicating an elongation of the distances upon mutation. white color represents low values which considered to have negligible impact on distances.

**Table S2:** The Difference contact map between Cas8 and nt-4 in the PAM sequence. Red color represents Negative values indicating a shortening in the distances upon mutation, whereas blue color represents positive values indicating an elongation of the distances upon mutation. white color represents low values which considered to have negligible impact on distances.

|  |  | CC | CG | Diff | TC | Diff | AC | Diff |
| --- | --- | --- | --- | --- | --- | --- | --- | --- |
| ASP51 | nt-4 | 8 | 7.94 | -0.06 | 8 | 0 | 8 | 0 |
| LYS52 | nt-4 | 8 | 5.08 | -2.92 | 8 | 0 | 8 | 0 |
| GLN67 | nt-4 | 8 | 2.96 | -5.04 | 8 | 0 | 8 | 0 |
| LYS68 | nt-4 | 8 | 2.84 | -5.16 | 8 | 0 | 8 | 0 |
| GLU72 | nt-4 | 8 | 6.42 | -1.58 | 8 | 0 | 8 | 0 |
| LYS73 | nt-4 | 6.69 | 8 | 1.31 | 8 | 1.31 | 8 | 1.31 |
| ASP129 | nt-4 | 5.15 | 6.99 | 1.84 | 6.1 | 0.95 | 5.29 | 0.14 |
| SER130 | nt-4 | 2.09 | 5.02 | 2.93 | 4.53 | 2.44 | 3.42 | 1.33 |
| LYS131 | nt-4 | 1.87 | 3.21 | 1.34 | 2.55 | 0.68 | 2.65 | 0.78 |
| TYR132 | nt-4 | 6.06 | 6.56 | 0.5 | 7.52 | 1.46 | 6.03 | -0.03 |
| VAL133 | nt-4 | 7.17 | 6.72 | -0.45 | 8 | 0.83 | 7 | -0.17 |
| ASN134 | nt-4 | 5.26 | 7.46 | 2.2 | 8 | 2.74 | 4.83 | -0.43 |
| LYS135 | nt-4 | 4.33 | 6.37 | 2.04 | 3.92 | -0.41 | 6.05 | 1.72 |
| ARG246 | nt-4 | 6.71 | 5.63 | -1.08 | 6.18 | -0.53 | 5.87 | -0.84 |
| THR248 | nt-4 | 3.93 | 6.08 | 2.15 | 6.95 | 3.02 | 5.55 | 1.62 |
| ASN249 | nt-4 | 5.81 | 7.14 | 1.33 | 5.75 | -0.06 | 5.71 | -0.1 |

**Table S3:** The Difference contact map between Cas8 and nt-53 in the PAM sequence. Red color represents Negative values indicating a shortening in the distances upon mutation, whereas blue color represents positive values indicating an elongation of the distances upon mutation. white color represents low values which considered to have negligible impact on distances.

|  |  | CC | CG | diff | TC | Diff | AC | Diff |
| --- | --- | --- | --- | --- | --- | --- | --- | --- |
| LYS91 | nt-53 | 8 | 8 | 0 | 7.82 | -0.18 | 7.87 | -0.13 |
| ILE95 | nt-53 | 8 | 8 | 0 | 3.89 | -4.11 | 4.9 | -3.1 |
| ARG96 | nt-53 | 8 | 8 | 0 | 4.8 | -3.2 | 4.15 | -3.85 |
| ASP129 | nt-53 | 8 | 8 | 0 | 7.81 | -0.19 | 7.02 | -0.98 |
| SER130 | nt-53 | 5.99 | 5.68 | -0.31 | 2.98 | -3.01 | 2.26 | -3.73 |
| LYS131 | nt-53 | 8 | 8 | 0 | 6.49 | -1.51 | 5.86 | -2.14 |
| TYR132 | nt-53 | 8 | 8 | 0 | 8 | 0 | 7.98 | -0.02 |
| VAL133 | nt-53 | 8 | 8 | 0 | 7.89 | -0.11 | 7.15 | -0.85 |
| PHE244 | nt-53 | 7.43 | 6.28 | -1.15 | 7.82 | 0.39 | 7.91 | 0.48 |
| SER245 | nt-53 | 5.44 | 2.47 | -2.97 | 4.27 | -1.17 | 5.25 | -0.19 |
| ARG246 | nt-53 | 1.74 | 1.98 | 0.24 | 3.04 | 1.3 | 2.71 | 0.97 |
| THR247 | nt-53 | 5.33 | 6.68 | 1.35 | 6.11 | 0.78 | 6.84 | 1.51 |
| THR248 | nt-53 | 2.27 | 4.86 | 2.59 | 2.67 | 0.4 | 2.57 | 0.3 |
| ASN249 | nt-53 | 1.92 | 2.5 | 0.58 | 2.01 | 0.09 | 1.76 | -0.16 |
| MET250 | nt-53 | 6.04 | 7.89 | 1.85 | 6.54 | 0.5 | 6.42 | 0.38 |
| GLY251 | nt-53 | 8 | 8 | 0 | 8 | 0 | 7.53 | -0.47 |
| MET255 | nt-53 | 8 | 8 | 0 | 7.98 | -0.02 | 7.93 | -0.07 |
| ALA457 | nt-53 | 8 | 8 | 0 | 7.62 | -0.38 | 8 | 0 |
| PHE459 | nt-53 | 8 | 8 | 0 | 6.62 | -1.38 | 8 | 0 |
| VAL460 | nt-53 | 8 | 8 | 0 | 5.93 | -2.07 | 5.45 | -2.55 |
| GLU461 | nt-53 | 8 | 8 | 0 | 8 | 0 | 7.09 | -0.91 |
| LYS462 | nt-53 | 7.32 | 1.73 | -5.59 | 2.73 | -4.59 | 1.8 | -5.52 |
| THR466 | nt-53 | 8 | 8 | 0 | 8 | 0 | 7.77 | -0.23 |
| ILE467 | nt-53 | 7.02 | 8 | 0.98 | 8 | 0.98 | 7.51 | 0.49 |
| SER468 | nt-53 | 4.98 | 4.68 | -0.3 | 8 | 3.02 | 4.59 | -0.39 |
| ALA469 | nt-53 | 2.69 | 5.71 | 3.02 | 8 | 5.31 | 4.28 | 1.59 |
| PRO470 | nt-53 | 5.88 | 5.02 | -0.86 | 2.41 | -3.47 | 2.41 | -3.47 |
| ALA471 | nt-53 | 5.45 | 2.54 | -2.91 | 2.09 | -3.36 | 3.63 | -1.82 |
| THR472 | nt-53 | 7.85 | 4.49 | -3.36 | 4.17 | -3.68 | 6.41 | -1.44 |
| ARG473 | nt-53 | 8 | 8 | 0 | 7.92 | -0.08 | 7.43 | -0.57 |

**Table S4:** The Difference contact map between Cas8 and nt-5 in the PAM sequence. Red color represents Negative values indicating a shortening in the distances upon mutation, whereas blue color represents positive values indicating an elongation of the distances upon mutation. white color represents low values which considered to have negligible impact on distances.

|  |  | CC | CG | Diff | TC | Diff | AC | Diff |
| --- | --- | --- | --- | --- | --- | --- | --- | --- |
| THR19 | nt-5 | 7.25 | 8 | 0.75 | 7.88 | 0.63 | 7.88 | 0.63 |
| THR20 | nt-5 | 8 | 8 | 0 | 7.92 | -0.08 | 8 | 0 |
| LYS23 | nt-5 | 4.08 | 8 | 3.92 | 6.05 | 1.97 | 4.02 | -0.06 |
| ARG27 | nt-5 | 5.42 | 8 | 2.58 | 7.26 | 1.84 | 6.15 | 0.73 |
| LEU49 | nt-5 | 6.11 | 8 | 1.89 | 8 | 1.89 | 5.98 | -0.13 |
| THR50 | nt-5 | 8 | 8 | 0 | 5.71 | -2.29 | 8 | 0 |
| ASP51 | nt-5 | 8 | 8 | 0 | 5.75 | -2.25 | 7.7 | -0.3 |
| LYS52 | nt-5 | 8 | 8 | 0 | 6.77 | -1.23 | 7.93 | -0.07 |
| THR53 | nt-5 | 8 | 8 | 0 | 8 | 0 | 6.43 | -1.57 |
| TRP74 | nt-5 | 7.46 | 7.35 | -0.11 | 7.73 | 0.27 | 6.29 | -1.17 |
| TYR126 | nt-5 | 8 | 8 | 0 | 7.79 | -0.21 | 7.66 | -0.34 |
| ASP129 | nt-5 | 5.89 | 5.82 | -0.07 | 5.93 | 0.04 | 6.07 | 0.18 |
| SER130 | nt-5 | 2.64 | 2.39 | -0.25 | 1.75 | -0.89 | 8 | 5.36 |
| LYS131 | nt-5 | 2.35 | 2.52 | 0.17 | 2 | -0.35 | 2.61 | 0.26 |
| TYR132 | nt-5 | 5.33 | 3.72 | -1.61 | 4.17 | -1.16 | 4.16 | -1.17 |
| VAL133 | nt-5 | 4.72 | 2.75 | -1.97 | 4.7 | -0.02 | 2.42 | -2.3 |
| ASN134 | nt-5 | 2.28 | 3.12 | 0.84 | 3.26 | 0.98 | 2.21 | -0.07 |
| LYS135 | nt-5 | 1.92 | 4.08 | 2.16 | 1.77 | -0.15 | 4.39 | 2.47 |
| SER136 | nt-5 | 7.39 | 6.56 | -0.83 | 7.13 | -0.26 | 7.8 | 0.41 |
| ARG246 | nt-5 | 6.21 | 2.34 | -3.87 | 4.5 | -1.71 | 2.69 | -3.52 |
| THR247 | nt-5 | 6.11 | 6.69 | 0.58 | 8 | 1.89 | 7.03 | 0.92 |
| THR248 | nt-5 | 1.57 | 1.94 | 0.37 | 2.97 | 1.4 | 1.78 | 0.21 |
| ASN249 | nt-5 | 5.24 | 2.08 | -3.16 | 4.22 | -1.02 | 4.56 | -0.68 |
| MET250 | nt-5 | 7.91 | 6.84 | -1.07 | 8 | 0.09 | 7.56 | -0.35 |
| GLY251 | nt-5 | 8 | 8 | 0 | 8 | 0 | 7.81 | -0.19 |
| VAL252 | nt-5 | 8 | 8 | 0 | 8 | 0 | 7.06 | -0.94 |
| MET255 | nt-5 | 6.71 | 4.22 | -2.49 | 5.04 | -1.67 | 3.81 | -2.9 |

## Notes

### Competing Interest Statement

The authors have declared no competing interest.

